# Natural reassortment of a segmented RNA arbovirus illustrates plasticity of phenotype in the arthropod vector and mammalian host *in vivo*

**DOI:** 10.1101/2021.08.09.455771

**Authors:** Christopher Sanders, Eva Veronesi, Paulina Rajko-Nenow, Peter Paul Clement Mertens, Carrie Batten, Simon Gubbins, Karin Darpel, Simon Carpenter

## Abstract

Segmented RNA viruses are a taxonomically diverse group of 11 families that can infect plant, wildlife, livestock and human hosts. A shared feature of these viruses is the ability to exchange genome segments during co-infection of a host by a process termed ‘reassortment’. Reassortment enables rapid evolutionary change, but in the case of segmented RNA viruses utilising an arthropod vector is set against the constraint of purifying selection and genetic bottlenecks imposed by replication in two evolutionarily distant hosts. In this study, we use an *in vivo* host: arbovirus: vector model to investigate the impact of reassortment on two phenotypic traits: vector competence and virulence in the host. Bluetongue virus (BTV) (*Reoviridae*) is the causative agent of bluetongue (BT), an economically important disease of domestic and wild ruminants and deer. The genome of BTV is comprised of 10 linear segments of dsRNA and the virus is transmitted between ruminants by *Culicoides* biting midges (Diptera: Ceratopogonidae). Five strains of BTV representing three serotypes (BTV-1, BTV-4 and BTV-8) were isolated from naturally infected ruminants in Europe and parental/reassortant lineage status assigned through full genome sequencing. Each strain was then assessed in parallel for the ability to infect *Culicoides* and to cause BT in sheep. Our results demonstrate that two reassortment strains, which themselves became established in the field, had obtained high replication ability in *C. sonorensis* from one of the parental virus strains which allowed inferences of the genome segments conferring this phenotypic trait.

**IMPORTANCE:** Reassortment between strains can lead to major shifts in the transmission parameters and virulence of segmented RNA viruses with consequences for spread, persistence and impact. The ability of these pathogens to change their phenotypes rapidly in response to selection pressure in new environments presents a major challenge in understanding factors driving emergence. Utilising a natural mammalian host-insect vector infection and transmission model, we demonstrated for the first time the genetic basis for a phenotypic trait of BTV within strains directly isolated from the field and, hence, selected and relevant for natural transmission.

## INTRODUCTION

Segmented RNA viruses include a diverse array of species classified across eleven taxonomic families that infect a wide range of hosts that include plants, animals, fungi, bacteria and marine protists (1). A key feature of segmented RNA viruses is their ability to exchange complete segments of RNA during coinfection of a single host cell by two or more virus strains, producing hybrid progeny. This form of recombination is termed ‘reassortment’. In the case of arboviruses (arthropod-borne viruses), selection of reassortant strains with advantageous phenotypic traits can occur through replication bottlenecks within both the hosts and biological vectors of the virus. When compared to genetic drift through mutation, reassortment can lead to more rapid changes in the phenotypic characteristics of progeny viruses and can lead to an increased transmissibility (2), increased pathogenicity (3, 4) and the potential for avirulent vaccine strains to revert to virulence in the field (5, 6).

Bluetongue virus (BTV) (*Reoviridae*) is the causative agent of bluetongue (BT), an economically important disease of domestic and wild ruminants (7). The virus is primarily spread between ruminants by *Culicoides* biting midges which act as biological vectors (8, 9). Severe clinical signs of bluetongue (BT) are most commonly observed in specific breeds of sheep (7) and are characterised by injury to the vascular and lymphatic endothelium. This can result in haemorrhage and vascular leakage that in acute cases result in fever, oedema, coronitis, oral and nasal erosion, cyanosis of the tongue and death (7, 10, 11). Cattle typically show only mild clinical signs of BT following infection, but are important reservoirs of the virus and recent outbreaks in naïve populations have documented more severe clinical signs in this species caused by specific BTV strains (12, 13).

Since the turn of the century, there has been an unprecedented shift in the epidemiology of BTV in Europe, involving the incursion of multiple strains into regions with no recorded history of transmission (8, 14). These epidemics have persisted in some cases and BTV has become endemic in several European countries, with major consequences for livestock production and trade (15, 16). Full-genome sequencing of BTV has demonstrated that reassortment occurs at a high frequency in the field (6, 17, 18) and this has been highlighted as a potential driver of virus emergence and spread in the region (6). BTV has a linear dsRNA genome consisting of 10 segments encoding seven structural (VP1-7) and at least four non-structural (NS1-4) proteins (19, 20).

Under experimental conditions, reassortment of genome segments between BTV strains during co-infection has been reported in the insect and ruminant hosts, where strains have been introduced into hosts, vectors or cell culture simultaneously (21–24). Arbovirus species and strain are also known to influence vector competence (25) and reassortment has been used to generate viral strains that express different levels of infection rate in *Culicoides* vectors in the laboratory (26). The proportion of a vector population able to become infected with BTV following oral exposure has been demonstrated to be under a combination of genetic and environmental control in *Culicoides* (9, 27, 28). To date, however, little is known regarding the impact of field-based reassortment of segmented RNA viruses on transmission by vectors and whether vector susceptibility to infection can be affected by this process.

Previous studies have also demonstrated variation in the clinical severity of BTV virus strains *in vivo* (7, 29–31), but the molecular basis for pathogenicity of BTV strains is poorly understood (32, 33). Pathogenicity and disease outcome appear to be highly complex and cannot be explained by the presence or absence of a single genome segment or specific combinations of segments (32–34). Reassortant bluetongue viruses generated from wild type and attenuated strains using reverse genetics demonstrated impacts on pathogenicity *in vitro* and *in vivo*, albeit with limited consistency between the different host systems used. To date, specific virulence characteristics could not be assigned to specific gene segments and, hence, no virulence markers for BTV strains exist (33, 35, 36). Field derived reassortant BTV strains may therefore demonstrate differential pathogenicity or disease outcome to that of their closely related strains or those which they share serotype. Studies with field-derived reassortant strains complement those carried out using BTV strains generated synthetically through reverse genetics, which have a far greater level of artificial selection through multiple cell passage and plaque purification (33, 37).

The aim of this study is to define phenotypic transmission characteristics of five BTV strains of three serotypes isolated from the Mediterranean Basin and Europe and to examine the impact of reassortment on virus phenotype. We identified a history of reassortment between these strains during their co-circulation in the field, which enabled the opportunity to examine the impact of this process on BTV phenotype in both host and vector. This represents the first fully comparative analysis of parental and reassortant strain phenotype under highly controlled conditions using a natural BTV-sheep-*Culicoides* transmission model.

## RESULTS

### Phylogenetic lineage of parental and reassortant strains can be traced

The genetic relationship between BTV strains used in the study (Table S1) was defined using sequence comparison between segments (Table S2), phylogenetic analyses (Fig S1) and a range of additional detection methods (Fig 1). Three strains were identified as parental (BTV-1 MOR2004/01, BTV-4 MOR2004/02 and BTV-8 NET2006/06), while two strains were identified as reassortant strains (BTV-4 MOR2009/07 and BTV-4 MOR2009/10) derived from the lineages of the three parental strains.

**Fig 1:**
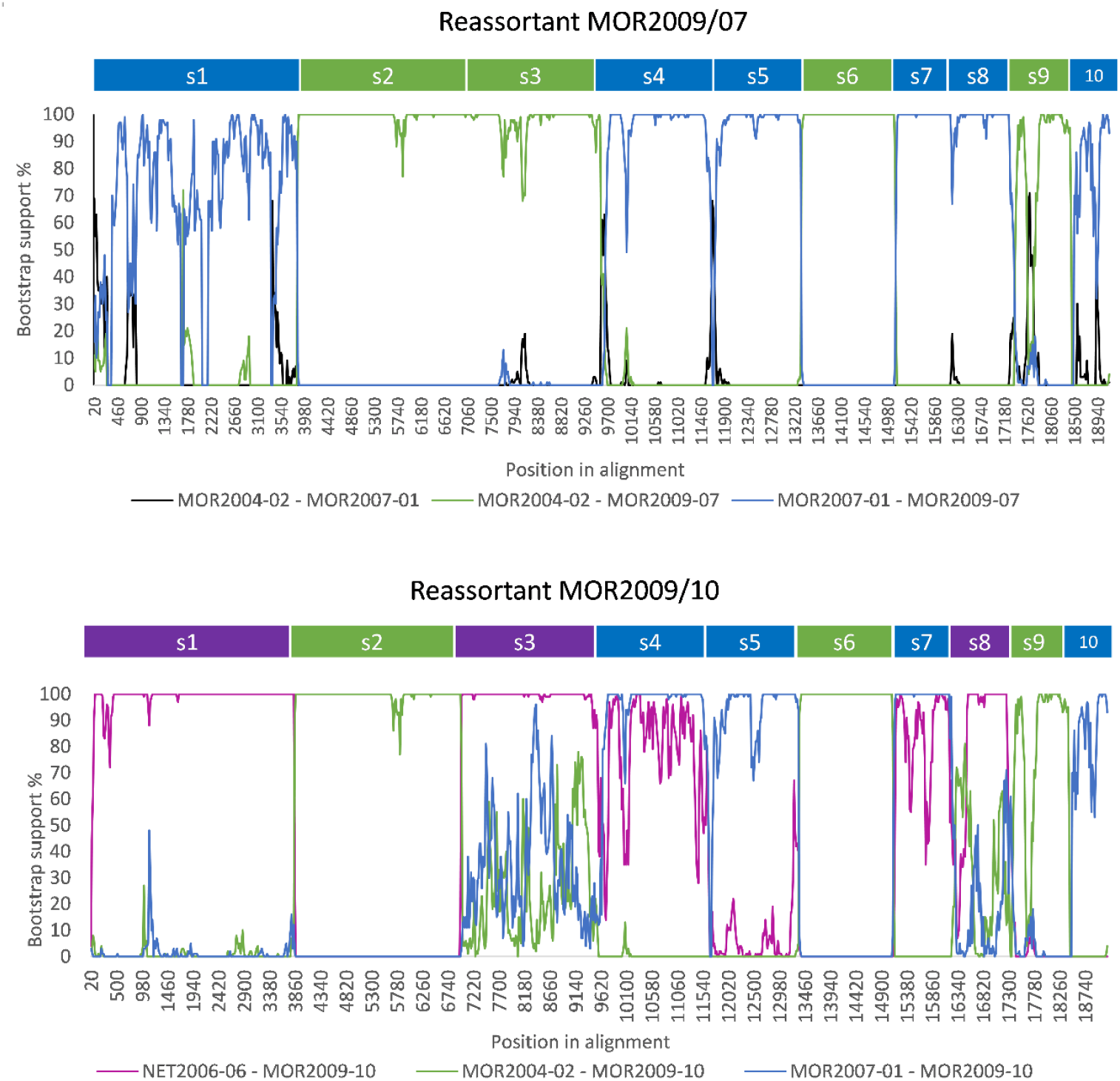
BOOTSCAN evidence for reassortment of BTV-4 MOR2009/07 (top) and BTV-4 MOR2009/10 (bottom).

The detection of the reassortant BTV-4 MOR2009/07 was supported by seven different detection methods within the recombination detection program: RDP (1.00×10^−10^), GENECONV (7.42×10^−110^), Bootscan (6.62×10^−131^), Maximum Chi Square (4.02×10^−28^), CHIMAERA (2.03×10^−29^), SISCAN (5.26×10^−31^) and 3SEQ (3.22×10^−100^). BTV-4 MOR2004/02 and BTV-1 MOR2007/01 sequences were identified as the major parent and the minor parent, respectively (Fig 1). The detection of the second reassortant (BTV-4 MOR2009/10) was supported by six different detection methods: GENECONV (4.55×10^−136^), Bootscan (4.64×10^− 4^), Maximum Chi Square (4.18×10^−27^), CHIMAERA (5.92×10^−36^), SISCAN (1.42×10^−37^) and 3SEQ (1.11×10^−15^). BTV-4 MOR2009/10 was identified as a triple reassortant, sharing segments from BTV-4 MOR2004/02 (Seg-2, 6 and 9), BTV-1 MOR2007/01 (Seg-4, 5, 7 and 10) and BTV-8 NET2006/06 (Seg-1, 3 and 8) strains (Fig 1, Table S2). It is possible that the BTV-4 MOR2009/10 emerged from a single cell co-infection with two strains (BTV-4 MOR2009/07 and BTV-8 NET2006/06) rather than simultaneous co-infection with three different BTV strains (BTV-1 MOR2007/01, BTV-4 MOR2004/02, and BTV-8 NET2006/06).

Two of the parental strains (BTV-1 MOR2007/01 and BTV-4 MOR2004/02) were isolated from samples collected in Morocco, while the third parental strain (BTV-8 NET2006/06) was isolated from a sample collected in the Netherlands. The field-derived reassortant strains, BTV-4 MOR2009/07 and BTV-4 MOR2009/10 were also isolated from samples collected in Morocco. The beginning and end recombination breakpoints (shown in Fig 1) corresponded with the segment position in the sequence alignment and are for Seg-1 (1-3944), Seg-2 (3945-6910) Seg-3 (6911-9682), Seg-4 (9683-11663), Seg-5 (11664-13441), Seg-6 (13442-15079), Seg-7 (15080-16235), Seg-8 (16236-17360), Seg-9 (17361-18411) and Seg-10 (18412-19233). Both reassortant strains had Seg-2 derived from BTV-4 MOR2004/02, and therefore both belonged to the same genotype/serotype (Fig S1).

As both the BTV-4 MOR2009/07 and BTV-4 MOR2009/10 reassortants were derived from the same parental strains, they were either identical or close to identical at the amino acid level in Seg-2 (99.8%), 4 (100%), 5 (100%), 6 (100%), 7 (100%), 9 (100%) and 10 (100%). However, BTV-4 MOR2009/07 and BTV-4 MOR2009/10 differed in the remaining segments: Seg-1 (99.4%), Seg-3 (99.9%), and Seg-8 (97.7%) (Table S2).

### Infection of sheep with BTV by infected *Culicoides* is highly efficient, independent of BTV strain

A total of 2762 *C. sonorensis* were intrathoracically inoculated (ITI) with the 5 strains of BTV, of which 1121 (40.6%) survived the incubation period and 486 (17.6%) successfully blood fed on sheep (Table S3). All sheep in all replicates were successfully infected with BTV despite ≤10 infected *C. sonorensis* taking a blood meal in 6 of 20 infection attempts and only one individual *C. sonorensis* taking a blood meal from a sheep on two occasions. All five strains of BTV were successfully transmitted on at least one occasion from the bites of 5 or fewer infected *C. sonorensis* during the trials (Fig 2; Fig 3).

**Fig 2.**
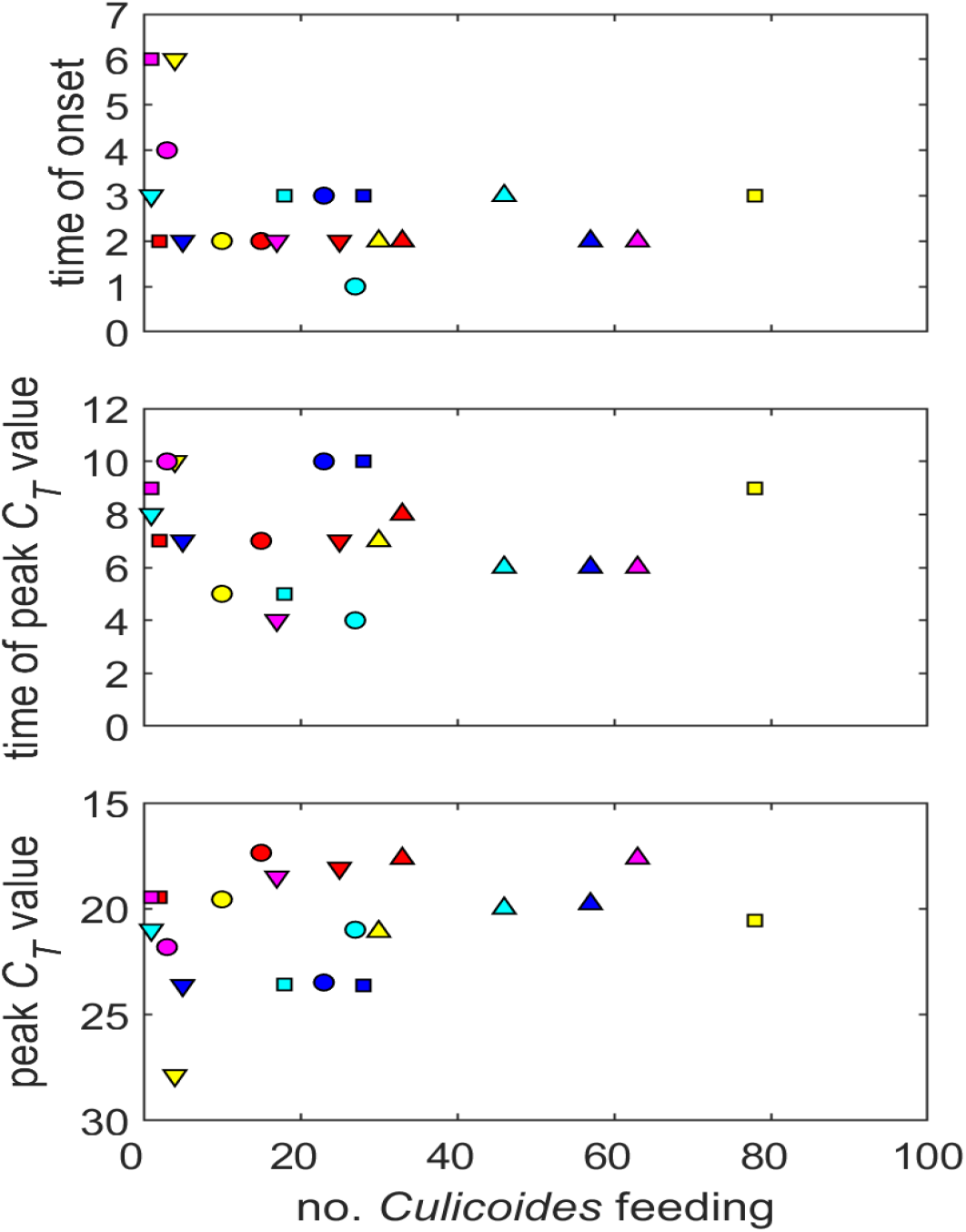
Time to onset of viraemia, time to peak viraemia and level of peak viraemia in sheep infected with different strains of bluetongue virus and their dependence on the number of inoculated *Culicoides* feeding to initiate the infection. Strains are: BTV-1 MOR2007/01 (red), BTV-4 MOR2004/02 (blue), BTV-8 NET2006/08 (cyan), BTV-4 MOR2009/07 (yellow) and BTV-4 MOR2009/10 (magenta). The different symbols indicate the different sheep infected with each strain.

**Fig 3.**
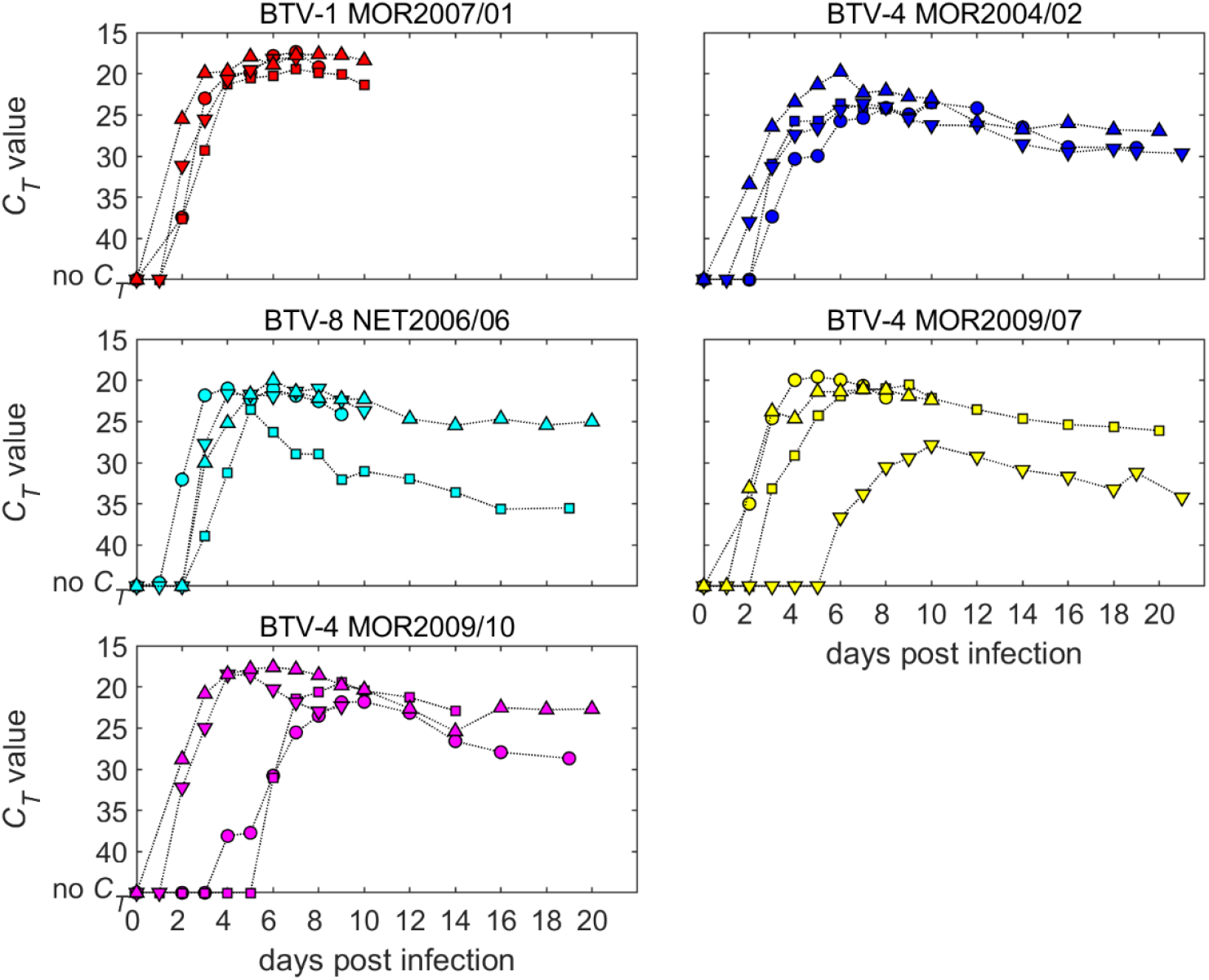
Viraemia in sheep infected with one of five strains of bluetongue virus. Each plot shows the changes in *C*_*T*_ value over time for individual sheep (indicated by symbols).

### Level of BTV viraemia in sheep varies with strain and number of infected *Culicoides* feeding

There was no significant effect of BTV strain (P=0.47) or number of *Culicoides* feeding (P=0.58) on the timing of peak viraemia (Fig 2; Fig 3). The level of peak viraemia differed significantly amongst BTV strains (P=0.009), with a higher level of RNA detected (i.e. lower *C*_*T*_ value) for BTV-1 MOR2007/01 compared with BTV-4 MOR2004/02 (P=0.02) and BTV-4 MOR2009/07 (P=0.02) (Fig 2; Fig 3). In addition, the level of peak viraemia increased significantly (i.e. the *C*_*T*_ value decreased) with the number of *Culicoides* feeding (P=0.02, b=-0.05, 95% confidence interval: −0.09 to −0.01) (Fig 2). At an individual replicate level, several sheep produced a late and slowly progressing viraemia (Fig 3). In these three cases, BTV RNA was not detected until 5 or 6 days post-infection. These cases involved BTV-4 MOR2009/07 transmitted by four *C. sonorensis* and two infections of BTV-4 MOR2009/10 by one and three biting midges, respectively (Table S3).

### Clinical disease in infected sheep varies with BTV strain and illustrates multi-factorial causation

Infection with BTV-1 MOR2007/01 caused acute clinical disease in all four exposed sheep necessitating euthanasia between 7-11 dpi for reaching pre-defined humane endpoints (Fig.4; Fig S2 Tables S4 and S5). Clinical disease in sheep infected with BTV-1 MOR2007/01 was highly uniform in manifestation, while all other BTV strains exhibited individual sheep variation in disease within infected cohorts. Three of the other strains produced acute clinical disease necessitating euthanasia in two sheep and milder, chronic disease in the other two (Fig 4; Fig S2;Table S5), while parental strain BTV-4 MOR2004/02 only led to acute disease in one sheep, which at post-mortem showed only mild pathological damage. Although the overall severity score of the parental strain, BTV-4 MOR2004/02 was comparable to those of the reassortment strains BTV-4 MOR2009/07 and BTV-4 MOR2009/10, the overall disease manifestation presented as more chronic, reflected in a higher clinical index score accumulation based on signs such reddening of eyes, nasal discharge and feet lesions (Figure 4; Fig S2; Table S5).

**Fig 4:**
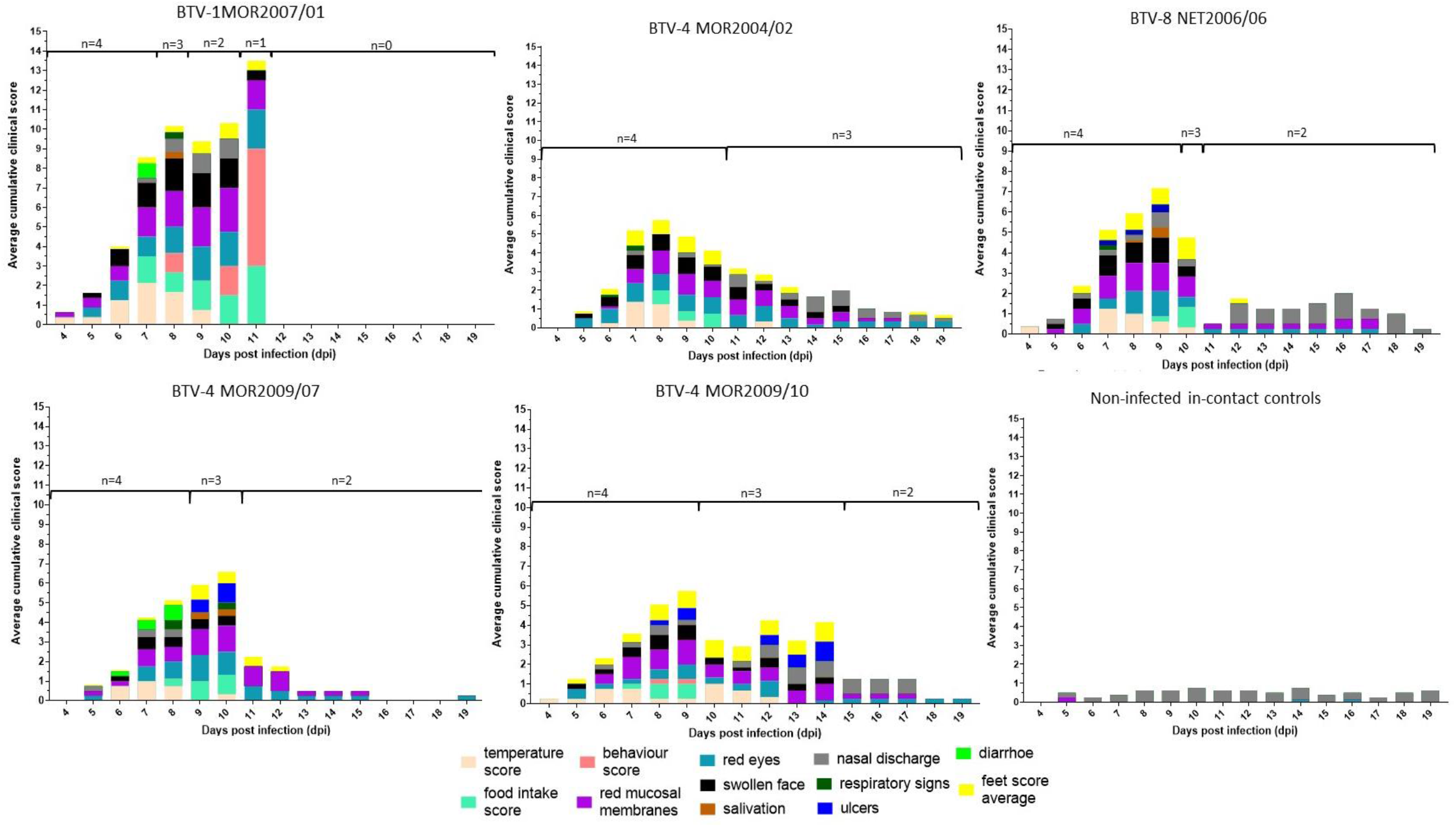
Daily accumulated average clinical score observed in sheep infected with 5 different BTV strains. Clinical scores were obtained across 12 different clinical signs routinely observed during BTV infection and visualised for each of the separate clinical sign for the time period of 4-19 dpi (see legend). The daily clinical scores recorded for each clinical sign were combined from all sheep within the group and then normalised to the sheep still present on the day to account for removal of sheep from groups at different days due to reaching predefined humane end points. Clinical scores for the uninfected in-contact controls were also recorded throughout to highlight potential non-specific clinical observations.

At post-mortem all four sheep infected with BTV-1 MOR2007/01 demonstrated systemic haemorrhages and oedema that was not only confined to the oral and nasal cavity, but also generalised through the subcutaneous layer and skeletal muscles (Figure S3). These lesions were less pronounced in BTV-8 NET2006/04 and further reduced in BTV-4 MOR2004/02. Three of the four acutely affected sheep infected with either the reassortant strains BTV-4 MOR2009/07 or BTV-4 MOR2009/10 developed overt ulceration, either of the oral or labial mucosa, or the muzzle, while only one sheep from the BTV-8 NET2006/04 cohort developed mild ulceration among all the parental strains (Figure S3). The acutely affected sheep infected with either reassortant strain also developed significant haemorrhages in the oral/nasal cavities that was slightly less serve than BTV-1 MOR2007/01, but exceeding those observed for BTV-4 MOR2004/02 and BTV-8 NET2006/04. All sheep which had exhibited mild or moderate clinical disease and survived to the study end at 19/20 dpi only demonstrated mild pathological signs of BTV infection at post-mortem, independent of the BTV strain.

### Susceptibility to infection in *Culicoides* varies significantly with BTV strain and can change rapidly via natural reassortment

A total of 5263 out of 6457 female *C. sonorensis* that were fed directly on viraemic sheep survived eight days of incubation across the four replicates (Fig 5; Table S6). In addition, a total of 2673 out of 4943 female *C. sonorensis* were successfully membrane-fed on matched blood from the viraemic sheep and survived the period of incubation (Table S6). The proportion of *Culicoides* in which BTV RNA was detected varied significantly between strains tested (Fig 5). *Culicoides sonorensis* was close to refractory to infection with parental strains BTV-1 MOR2007/01 (2.4%) and BTV-8 NET2006/04 (0.4%), but highly susceptible to infection with the third parental strain BTV-4 MOR2004/02 (21.6%). Both BTV-4 MOR2009/07 (28.3%) and BTV-4 MOR2009/10 (45.0%) retained this infectivity following reassortment. The effect of blood-feeding-route (sheep vs membrane-based system) differed amongst the strains, but broadly recapitulated the clear difference in infectivity between BTV-1 MOR2007/01 and BTV-8 NET2006/04 in comparison to the three other BTV strains (Fig 5; Fig 6; Table S7). In BTV-4 MOR2004/02, membrane feeding was associated with decreased viral replication in the insect vector compared with feeding on an infected sheep, while for BTV-4 MOR2009/07 it was associated with an increase. For the other three strains, there was no significant difference in vector infection efficiency between the two feeding routes (Fig 6; Table S7).

**Fig 5.**
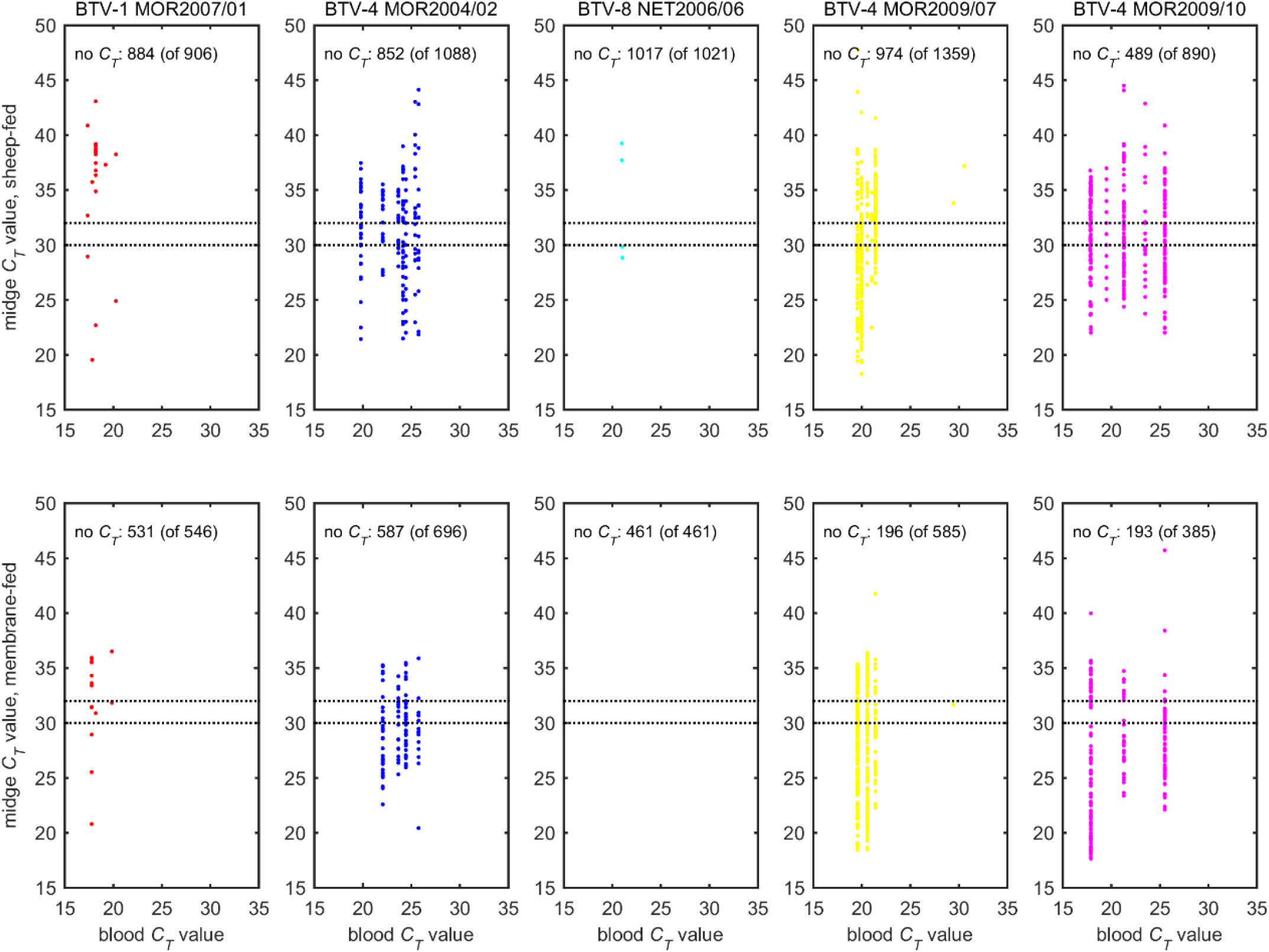
Observed *C*_*T*_ values for five strains of bluetongue virus in *Culicoides sonorensis* following blood feeding and incubation for eight days at 25 °C. Each plot shows the dependence of the *C*_*T*_ value on feeding route (sheep-fed: top row; membrane-fed: bottom row) and the *C*_*T*_ value of the infected blood meal. The numbers at the top of each plot indicate the number of midges with no *C*_*T*_ value after incubation and the total number of *C. sonorensis* tested. The dotted lines indicate *C*_*T*_ values which correspond to a possible (*C*_*T*_<32) and probable (*C*_*T*_<30) fully transmissible infection in a previous study of BTV infection in *Culicoides* (Veronesi *et al*. 2013).

**Fig 6.**
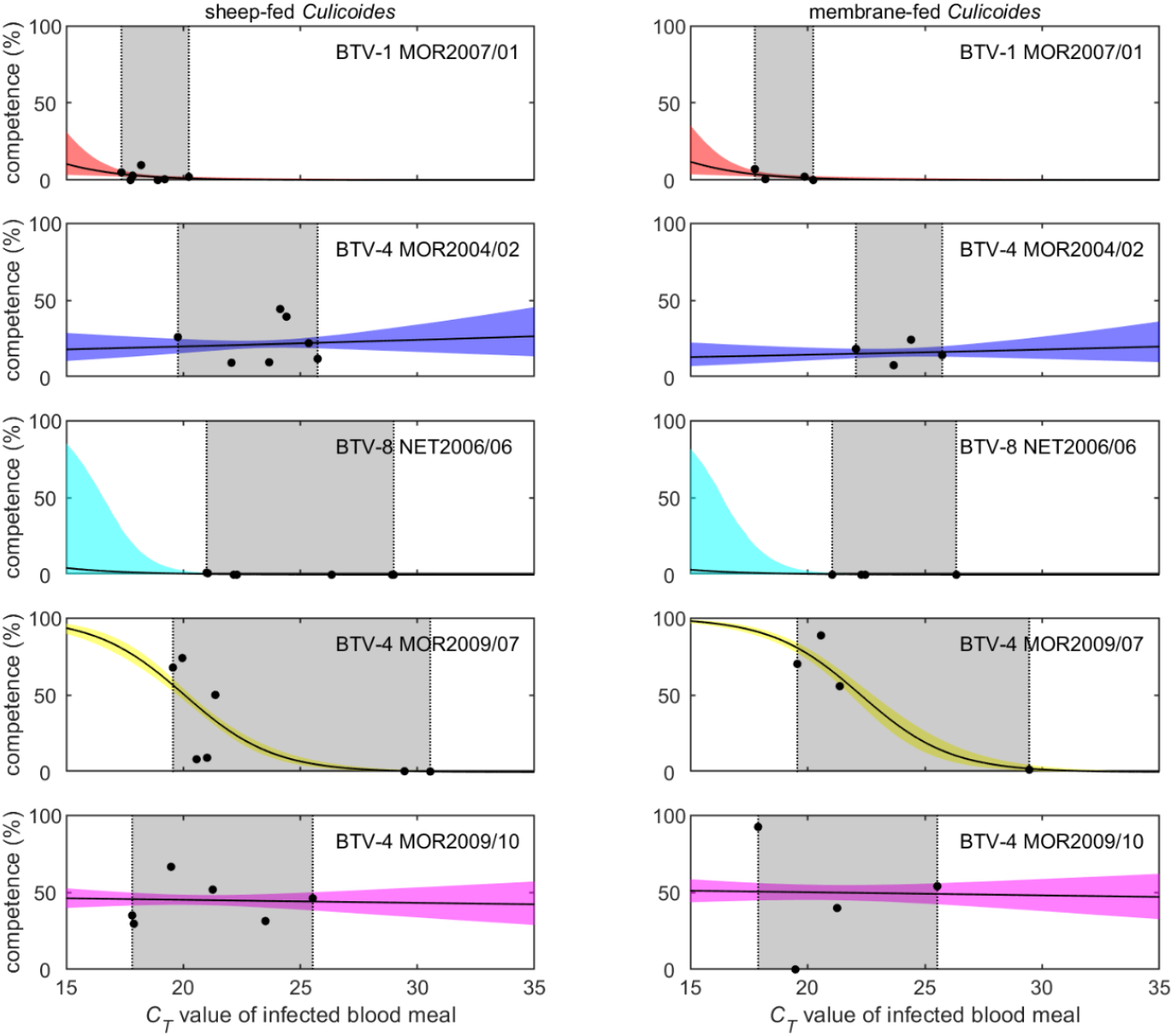
Vector competence of *Culicoides sonorensis* for five strains of bluetongue virus and its dependence on feeding route and blood meal *C*_*T*_ value. Each plot shows the posterior median (black line) and 95% credible interval (shading) for the vector competence. The black circles are the observed proportions of infected midges at each *C*_*T*_ value and the grey shaded area represents the range of *C*_*T*_ values for the infected blood meals.

## DISCUSSION

This is the first study to demonstrate the impact of natural reassortment on biological transmission of a segmented RNA arbovirus between ruminant and arthropod hosts. Using a highly repeatable and manipulatable *Culicoides*-BTV-sheep model and full genome sequencing data, we show that inheritance of phenotype during reassortment was explanatory of vector susceptibility to infection, while virus pathogenicity following reassortment was unpredictable. These observations have fundamental consequences both for understanding selective pressures operating on reassortment in the field and in the ability to predict the potential for rapid shifts in virus phenotype within regions where virus strains circulate.

While tracing a direct lineage between reassortant strains of BTV is challenging, our sequencing data analyses suggested a high degree of similarity in segments inherited from parental strains, indicating negligible genetic drift within segment sequences. This inference was aided by a clear timeline in the isolation of strains used for the study and the use of parallel, rather than sequential, comparison of both clinical signs in sheep and transmission rates to *Culicoides* that underpinned phenotype definition. Comparisons of both clinical severity of BTV strain infection in hosts (29, 31, 38) and transmission to vectors (39) have been carried out previously. However, this is the first study to link both these parameters with viruses of known reassortment lineage.

Transmission of parental and reassortant strains of BTV from ITI *Culicoides* to naïve sheep was highly efficient across all five strains of BTV, confirming previous studies (40, 41). Interestingly, severity of clinical infection in sheep was not broadly correlated to the number of infective bites received, but, where very few fully infected *C. sonorensis* fed, this occasionally led to a late and slowly progressing viraemia. During the 2006/7 epidemic of BTV-8 in northern Europe (15), initial outbreaks of disease were classified as being mild which was initially attributed to reduced exposure to *Culicoides* biting as the incursion began in late summer (42). Our study suggests a decline in populations of *Culicoides* in autumn is unlikely to change the severity of report cases and clinical surveillance should still be effective (43, 44).

Rates of viral replication in *C. sonorensis* for four of the five BTV strains were comparable between membrane feeding on viraemic sheep blood and feeding directly from the sheep itself. The exception to this was the BTV-4 MOR2009/07 strain which produced a significantly higher rate of vector competence in membrane fed individuals. It is unclear at present why this strain-specific variation occurred as blood meal titres across the testing were similar. Moreover, this result was influenced by a high feeding rate on a single sheep and the results should therefore be treated with caution. One potential explanation could be that as the RNA quantities were estimated in blood taken from a superficial vein, whereas local virus replication in the skin might lead to local differences from systemic titres (45), leading to variation in vector competence when assessed. More broadly, however, it has been demonstrated here that the systemic level of BTV infection even when determined in blood samples taken at the jugular vein is a valid predictor of vector infection.

No specific pattern was demonstrated in gene segment inheritance for the severity of clinical disease within infected sheep, in part due to the wide variation in observations within cohorts. This finding was consistent with previous studies attempting to elucidate the genetic basis of BTV strain virulence, which also could not identify specific genome segments determining virulence in ruminants (32, 33). It is interesting to note that virulence still presented as a multi-segmented trait in this study, even when low-passage field isolates were transmitted to the sheep host via the most natural route of blood-feeding infected *C. sonorensis*. Infection with BTV-1 MOR 2007/01 did, however, result in a uniform manifestation of what would be considered to be classical clinical BT as described from field observations (7). In contrast all other strains led to a far more typical intra-cohort variation in BT severity using the clinical scoring approach. BTV-8 NET 2006/06 was somewhat less severe than expected from previous field reports (12, 13), although this is consistent with previous experiments (16, 31). Anecdotal observation had previously reported that BTV-4 strains increased in severity throughout time following co-circulation with BTV-1 and BTV-8. While the acute clinical presentation of sheep infected with the early parental BTV-4 MOR2004/02 strain was milder in comparison to the two BTV-4 reassortant strains MOR2009/07 and MOR2009/10, this was not reflected in the overall average severity score (Figure 3, Figure S2 and Table S5), partly through the more chronic presentation of disease in sheep infected with BTV-4 MOR2004/02 still leading to an overall significant clinical impact. Although the pathological damage caused by BTV-4 MOR2004/02 was clearly less severe, this is also compounded by the fact that sheep euthanised during peak clinical disease presented as vastly more acute during post-mortem examination compared to those examined later at study end. It is evident, however, that neither of the two BTV-4 reassortant strains inherited the full virulence of its primary parental strain BTV-1 MOR2007/01 despite six (BTV-4 MOR2009/07) or four (BTV-4 MOR2009/10) segments originating from this viral lineage.

Infection rates of *Culicoides* fed on viraemic sheep, in contrast to virus virulence, varied consistently and provided a clear marker of BTV strain phenotype during the studies. *Culicoides sonorensis* was almost entirely refractory to two of the parental strains (BTV-1 MOR 2007/01 and BTV-8 NET2006/06), while the third gave among the highest rates of infection recorded (BTV-4 MOR2004/02). This extremely high rate of infection was retained within both of the reassortant progeny (BTV-4 MOR2009/07 and BTV-4 MOR2009/10). To investigate the impact of amino acid variations of each viral protein (encoded by the specific genome segments) on vector competence, we investigated the BTV genomes for amino acid positions that were identical for all three BTV-4 strains, but different for both BTV-1 and BTV-8 parental strains.

Seg-7 was the most conserved across all five BTV isolates. Four out of five strains shared 100% amino acid similarity in this segment, whereas BTV-4 MOR2004/02 differed by one very conserved amino acid substation (Table S2). Therefore Seg-7 does not influence the infection rate of these five BTV strains in *C. sonorensis*. Although there were some amino acid variations in proteins encoded by segments −1 (99.0-99.7%), −3 (99.6-100%), −4 (98.4-100%), −8 (97.7-100%), and −10 (94.7-100%) (Table S2), no single position was identified that would be identical for all three BTV-4 strains, but different for both BTV-1 and BTV-8. Therefore, these segments also seem to have limited impact on the infection rate of BTV in *C. sonorensis* observed for these strains.

Seg-2, 6 and 9 of the BTV genome, encoding for VP2, VP5 and VP6 respectively, were shared across the three BTV-4 strains that exhibited a high rate of vector infection. For segment-9, four amino acids substitutions have been identified in all three BTV-4 strains, but not in BTV-1 MOR2007/01 and BTV-8 NET2006/06. Of these four amino acids substitutions, three were classified as conservative and only one as radical (position 95, Glycine to Arginine). It is unclear if this single amino acid substitution could have any effect on vector competence, but this could be further investigated in the future by generating reverse-engineered mono-reassortant strains.

Most notably, numerous amino acid substitutions have been identified in Seg-2 and -6 between the three BTV-4 strains and the other two parental strains BTV-1 MOR2007/01 and BTV-8 NET2006/06. It therefore seems highly like that VP2 and/or VP5 proteins derived from the parental BTV-4 MOR2004/02 play a key role in determining the vector competence, either individually or in combination, for the BTV strains used in this study. Viral proteins including VP2 from the atypical BTV 26 have been shown to restrict infection and therefore transmission in *Culicoides (46, 47)*. The BTV outer capsid protein VP2 is known to be highly variable, containing the epitopes to the host’s neutralising antibody response, thereby determining the strain serotype (48). Both outer capsid proteins VP2 and VP5 are responsible for cell entry (48), whilst VP5 also plays a role in the penetration of mammalian and insect cells (49) during the release of the core particle from the endosome. The affinity of VP2 for erythrocyte glycoprotein may facilitate transmission from mammalian blood to the vector (48). Furthermore, VP2 is cleaved by proteases in the saliva of competent vector *Culicoides* resulting in increased infectivity to *Culicoides* derived cells (50). The role of VP2 in the binding, entry and infection of *Culicoides* mesoenteron gut cells, the barrier to infection (51), is as yet undetermined.

This study has demonstrated that rapid phenotypic changes in co-circulating strains of BTV can occur, driven by reassortment, even in the presence of purifying selection and genetic bottlenecks imposed by utilising only distantly related hosts and vectors. From a policy perspective, the emergence of reassortant strains of BTV possess additional complexity both in the decision to implement vaccination campaigns and in the likelihood of spread in local vector populations. Currently, low pathogenicity strains of BTV are often allowed to spread where their impact is perceived as being less damaging than the cost of vaccination (15). In addition, the factors underlying the spread of BTV strains between regions dominated by different species of *Culicoides* are poorly understood and reassortment could allow these barriers to be overcome.

A future step is to examine the genetic drivers of this process by sequencing of strains exhibiting specific phenotypic characteristics in hosts and vectors. The application of next generation sequencing will be informative in understanding how the virus populations interacting in the process of reassortment are sustained and selected in both the ruminant and insect host. Studies could also be extended to examine tropism of virus communities in both the ruminant and insect host with a view to understanding dissemination. Subsequent testing of viruses produced using reverse genetics and informed by sequencing could also elucidate the genomic basis of a range of phenotypic responses including transmission probability, temperature limits to replication and pathogenicity in the ruminant host.

## MATERIALS AND METHODS

### Full genome sequencing of BTV strains

Full genome sequences of BTV strains were obtained by Sanger sequencing (BTV-4 MOR2004/02, BTV-1 MOR2007/01, BTV-4 MOR2009/07 and BTV-4 MOR2009/10), with the exception of BTV-8 NET2006/06 which was additionally resequenced using High Throughput Sequencing (HTS) (BTV-8 NET2006/06). Sanger sequencing was carried out as previously described (6). For HTS, total RNA was extracted from cell culture pellets using TRIzol Reagent (Life Technologies, Paisley, UK) and eluted in 100 µL of nuclease free water (Sigma-aldrich, Gillingham, UK). One microlitre of RNase T1 enzyme was added into each tube and incubated at 37 °C for 30 min in a thermocycler to remove ssRNA. DsRNA was purified using the RNA Clean and Concentrator kit (Zymo, Irvine, CA, USA) according to the manufacturer’s recommendations. The purified dsRNA (8 µL) was denatured by heating at 95 °C for 5 min and the first cDNA strand synthesised using SuperScript III RT (Life Technologies, Paisley, UK) while the second strand was synthesised using NEBNext (New England BioLabs, Hitchin, UK) according to the manufacturers’ instructions. Double stranded cDNA was quantified using the Qubit dsDNA HS Assay kit (Life Technologies) and then adjusted to 0.2 ng µL^−1^ with 10 mM Tris-HCl, pH 8.0 buffer. Library preparation was performed using the Nextera XT library preparation kit, and paired end read sequencing (2 x150bp) was performed using MiSeq platform and reagent kit v2 (Illumina, San Diego, CA, USA). For sequences obtained by HTS a pre-alignment quality check was performed using the FASTQC program v0.11.8 and the Trim Galore script (52) was used for quality and adapter trimming of FASTQ files along with removal of short sequences (<50 bp). Subsequently, reads were mapped to a reference sequence using the BWA-MEM tool (53) and then the DiversiTools software [http://josephhughes.github.io/DiversiTools/; accessed 12/10/2019] was used to generate the consensus sequence. Finally, the consensus sequence was used as a reference sequence to increase the number of BTV reads mapped to the reference, and the final consensus sequence was saved and used for further analysis. Genbank accession numbers for the BTV-8 NET2006/06 strain are MW159097-MW159106.

### Reassortment analysis

Reassortment analysis was performed using Recombinant Detection Program version 4.95 (RDP4) (54) under default settings. Previously sequenced and published strains were retrieved from GenBank: BTV-1 MOR2007/01 (KP820890, KP821010, KP821132, KP821252, KP821372, KP821492, KP821614, KP821734, KP821855, KP821975), BTV-4 MOR2004/02 (KP820941, KP821061, KP821183, KP821303, KP821423, KP821543, KP821665, KP821785, KP821905, KP822026), BTV-4 MOR2009/07 (KP820942, KP821062, KP821184, KP821304, KP821424, KP821544, KP821666, KP821786, KP821906, KP822027) and BTV-4 MOR2009/20 (KP820945, KP821065, KP821187, KP821307, KP821427, KP821547, KP821669, KP821789, KP821909, KP822030). Multiple sequence alignment was performed separately for each viral segment using the Muscle algorithm in MEGA6 program (55), coding regions were aligned on the amino acid level. Then, individual alignment files were concatenated using SequenceMatrix software (56) and NEXUS files were used for RDP analysis. The detection of potential recombination events was performed with the following methods: RDP, GENECONV, Maximum Chi Square, CHIMAERA, BOOTSCANing, Sister Scanning (SISCAN) and 3SEQ. Strains BTV-4 MOR2004/02, BTV-1 MOR2007/01 and BTV-8 NET2006/06 were considered as parental sequences based on their year of detection, BTV-4 MOR2009/07 and BTV-4 MOR2009/10 were investigated as potential reassortant strains.

### Standardization of viral inoculum

All isolates used were passaged once on *C. sonorensis* derived KC cells to generate infectious tissue culture supernatant (TCS) that was derived in the same cell culture system (Table S1). Briefly a T175 flask of KC cells in suspension of growth media (Schneider’s insect cell media (Sigma-Aldrich, Dorset, UK); 10% Gibco^®^ heat-inactivated fetal calf serum (ThermoFisher Scientific, MA, USA) and 1% Penicillin/Streptomycin (Sigma-Aldrich, Dorset, UK)) was inoculated with the stock virus from the Orbivirus reference collection and incubated for 7 days at 25°C. Following incubation, cells and supernatants were collected and cells pelleted through centrifugation for 10 min at 1000g and 4°C.

The supernatants of each BTV strain were titrated on KC cells using a 96-well end-point dilution microtitration assay. 10-fold serial dilutions were added to the titration plates containing 1×10^5 KC cells/well in suspensions of growth media. Titration plates were incubated for 7 days at 25°C at which point supernatants were removed and cells were fixed using 4% paraformaldehyde for 30-45 minutes, washed 3 x with PBS and stored under 100µl PBS/well at 4°C. As KC cells do not develop cytopathic effects, BTV infection of each well was determined through visualisation of BTV antigen by immunofluorescence microscopy. Cells were permeabilised with 0.2% Trition for 15 min followed by incubation with anti-BTV guinea pig hyperimmune serum (ORAB279) at 1:2000 in PBS-0.5%BSA for 1h at room temperature (RT). Following another 3 washes with PBS plates were incubated for 1h with secondary goat anti-guinea pig-AlexaFluor488 (Invitrogen, UK) at 1:500 in PBS-0.5%BSA. After 3x final PBS washes plates were examined under the fluorescence microscope and the presence or absence of fluorescence was recorded for each well.

The final infectious titre for each virus was calculated according to the Spearman-Karber method. Serotype specificity and the absence of cross-contamination between the different virus inocula was confirmed by serotype specific qRT-PCR (57). Viral inoculum characteristics are summarised in Table S1.

### Infection of *Culicoides* with BTV strains

Colony-derived adults of *Culicoides sonorensis* Wirth & Jones, a BTV vector in North America, were used in the study (58). Maintenance was as described previously (59), with the exception that the colony was sustained using a Hemotek artificial feeder and horse blood from a commercial supplier (TCS Bioscience, Botolph Claydon, UK). *Culicoides sonorensis* were intrathoracically (IT) inoculated with 0.2 µl BTV tissue culture supernatant at a standardised titre of 5.75 log_10_TCID_50_ for each BTV strain using a pulled glass capillary needle (Narishige, Japan) and a Nanoject II microinjector (Drummond Scientific, PA, USA). Between 50 and 200 *Culicoides* were inoculated for each virus strain in each experimental replicate and were subsequently incubated at 25°C (Binder, Tuttlingen, Germany) in card pillboxes with mesh screen for five to six days (60). *Culicoides* were given access to 10% sucrose solution on cotton wool that was replenished daily throughout this period.

### Animal experiment

This animal experiment was carried out in accordance with the UK Animal Scientific Procedure Act (ASPA) 1986 which transposes European Directive 2010/63/EU into UK national law. All animal procedures carried out were reviewed and approved by the Animal Welfare and Ethics Review Board at the Pirbright Institute and conducted in compliance with the relevant project licences granted by the UK Home Office. The study was conducted under high biological containment at the Pirbright Institute in 4 sequential replicates, each consisting of 6 sheep with one sheep in each replicate exposed to one parental or reassortant BTV strain and an associated uninfected control animal (24 sheep total). The sheep used were British mule crosses. All sheep were tested to confirm the absence of anti-BTV antibodies by the UK BTV reference laboratory prior to arrival at The Pirbright Institute using a commercial competitive anti-VP7 antibody ELISA (IDVet, Montpellier, France). All animals were allowed to acclimatise to the new facilities for 6-8 days before onset of any procedures and were fed twice a day with grain pellets and with ad libitum access to hay and water throughout the experiment (10).

#### Infection of sheep

Pots containing ITI *Culicoides* were placed on the inner thigh of a restrained sheep to feed for a period of 10 minutes. The strain of BTV used to infect each sheep and the sheep identification number were recorded. The *Culicoides* were anaesthetised and examined under a light stereomicroscope for evidence of blood feeding. Engorged, or partially engorged, individuals were placed in a microtube containing in 100 µl of either Schneider’s media (Experimental Replicates 1 and 2) or RPMI media (Experimental Replicates 3 and 4) to which 2% Penicillin/Streptomycin and Amphotericin B (Sigma-Aldrich, Dorset, UK) had been added. Samples were homogenised for two cycles of 30 seconds at 25Hz with a 3mm stainless steel bead in a Tissuelyser™ (Qiagen, Manchester, UK). The homogenates were then diluted to a total volume of 1000 µl with RPMI media (Sigma-Aldrich, Dorset, UK) and 2% antibiotics, the stainless steel bead removed and samples were then vortexed at 13,000 rpm for 10 minutes and stored at +4°C. 50 µl of homogenate was then used for nucleic acid extraction and analysis by rtRT-PCR to determine if the *Culicoides* had supported a disseminated infection and that transmission of BTV to the sheep was likely to have occurred.

#### Monitoring of BTV infection in sheep

Blood samples were taken from the jugular vein on the day before the experiment began (day 0) and then on days 1, 2, 3, 4, 5, 6, 7, 8, 9, 10, 12, 14, 16, 18 and 19/20 following infection. Clinical signs of disease and rectal temperature were recorded for each animal daily. A post-mortem of all sheep was carried out following euthanasia at the study end (19/20 dpi) or when individuals reached a humane endpoint for the protocol.

#### Calculation of clinical scores

A cumulative daily clinical score was determined for each cohort of four sheep infected with a single BTV strain and the four control animals (Table S4). To account for sheep euthanized due to exceeding clinical endpoints, daily accumulated clinical scores for each cohort were divided by the number of sheep present. A total severity score (Figure S2 and Table S5) for each virus was obtained by dividing the overall accumulated total clinical score from all sheep by the combined days the entire cohort of four individuals stayed alive between 4-19 dpi (period for which any clinical signs were recorded).

### Transmission of BTV strains from viraemic sheep to *Culicoides*

The quantity of BTV RNA in blood samples taken from sheep in the first ten days post-infection was tested on the day of collection. Preliminary data suggested that a blood sample with a sqPCR *C*_*T*_ value of less than 25 indicated that BTV infection would reach peak viraemia in the sheep within the following two days and this was used as a guide to time blood-feeding of *C. sonorensis* (40). *C. sonorensis* were allowed to feed on sheep in card boxes through a fine mesh lid at a density of approximately 250 individuals. The boxes were placed on the inner thigh of a restrained sheep for a period of 10 minutes and two boxes of *Culicoides* were fed on each sheep in each replicate. Replete female *C. sonorensis* were selected under light CO_2_ anaesthesia using a stereomicroscope. Blood-fed *C. sonorensis* were placed in new card pill boxes and incubated at 25°C with access to a 10% sucrose solution on cotton wool pads that were replenished daily. After an eight day incubation period post-feeding, all surviving female *Culicoides* were selected under CO_2_ anaesthesia and placed individually in microtubes (Qiagen) containing 100µl Schneider’s media (Experimental Replicates 1 and 2) or RPMI (ThermoFisher Scientific) (Experimental Replicates 3 and 4) for homogenisation as described previously. Homogenates were then diluted with 9000µl of either Schneider’s media or RPMI to a total of 1 ml, sealed and stored at +4°C.

### Comparison of feeding methods on infection rate in *Culicoides*

Natural blood feeding of *Culicoides* on viraemic sheep was complemented by artificial feeding of *C. sonorensis* on sheep blood taken on the matched day of peak viraemia for each strain in each replicate. Identical boxes of *C. sonorensis* were fed on sheep blood collected the same day as direct feeding. *Culicoides* were fed through a Parafilm™ (SigmaAldrich, Dorset, UK) membrane over a reservoir (Hemotek Ltd, UK) filled with 3 ml of viraemic sheep blood and maintained at 37°C. Following a period of 30 mins exposure to the blood, replete females were selected out under CO_2_ anaesthesia and treated as described for the *Culicoides* fed directly on sheep. Any individual *Culicoides* sample with a *C*_*T*_ value <40 when tested by PCR was considered BTV positive.

### Nucleic acid extraction and semi-quantitative rtPCR

Nucleic acid extraction throughout the studies was carried out using extraction robots. In replicates 1 and 2 a Universal extraction robot (Qiagen, Germany) and associated nucleic acid extraction (Qiagen, Germany) kits were used. In replicate 3 and 4, a Kingfisher Flex extraction robot (Thermo Fisher Scientific) and associated kits (MagVet™ Universal Isolation Kit) were used. Validation of the Kingfisher Flex against the Universal extraction robot demonstrated variation between machines was similar to variation observed between repeat runs on the same equipment (+/-0.5 *C*_*T*_) (data not shown). Analysis by sqPCR for all samples used a BTV group specific PCR against segment 1 of the virus genome (61). For replicates 1 and 2 a Strategene Mx3005P PCR machine (Agilent, USA) was used, in replicates 3 and 4, a Life Systems 7500 Fast PCR machine (Thermo Fisher Scientific, UK) was used. In validation between these machines a similar variation was observed to that between repeat runs on the same equipment (+/-0.5 *C*_*T*_). Confirmation of BTV serotype of viral stocks were carried out using a serotype specific rRT-PCR assay (57).

In assessments of *Culicoides* infection rates following blood feeding on sheep or through the membrane-based system, 50µl of pooled homogenate from 8 individually homogenised *Culicoides* was added to a microtube. From this 400µl pool, nucleic acid was extracted from a 50µl sample and viral RNA assessed by sqPCR as described above. Individuals contributing to a pooled sample found to contain BTV RNA (≤40 Ct), were subsequently tested individually by sqPCR.

### Statistical analysis

The effect of strain and number of *Culicoides* feeding at infection on timing and magnitude of peak viraemia (i.e. the lowest *C*_*T*_ value) was assessed using linear models with time of peak viraemia or *C*_*T*_ value at peak viraemia as the response variable and strain and number of *Culicoides* feeding and an interaction between them as explanatory variables. Model selection proceeded by stepwise deletion of non-significant (*P*>0.05) terms as judged by *F*-tests.

The vector competence (i.e. the probability of a *Culicoides* midge having a *C*_*T*_ value) after feeding on blood infected with strain *s* by route *f* (either on a sheep (*f*=0) or via a membrane (*f*=1)) when the *C*_*T*_ value of the infected blood meal is *c* is given by;

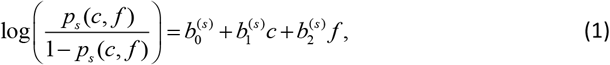

where the *b*_*i*_s are strain-specific parameters. Differences amongst strains were incorporated by assuming hierarchical structure for the model parameters, so that the parameters for strain *s* are drawn from higher-order normal distributions, namely,

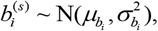

where the *μ*s and *σ*s are higher-order parameters.

Parameters were estimated in a Bayesian framework. A Bernoulli likelihood was used for the infection status of each *Culicoides* midge with probability of infection given by equation (1). Priors for the strain-specific parameters were given by the higher-order distributions, while non-informative priors were assumed for the higher-order parameters (diffuse normal for the *μ*s and diffuse gamma for the *σ*s). The methods were implemented in OpenBUGS (version 3.2.3;http://www.openbugs.info). Two chains each of 10,000 iterations were generated (with the first 2,000 iterations discarded to allow for burn-in of the chains). Convergence of the chains was monitored visually and using the Gelman-Rubin statistic in OpenBUGS.

## ACKNOWLEDGEMENTS

Funding for this project was provided by Defra grant SE:2618, with additional support provided by the Biotechnology and Biological Sciences Research Council (BBSRC) grants BBS/E/I/00001701, BBS/E/I/00001717, BBS/E/I/00001728, BBS/E/I/00007030, BBS/E/I/00007036, BBS/E/I/00007037 and BBS/E/I/00007039. We would like to that all the animal technicians involved in this study for their dedication and invaluable experience and Eric Denison for supply of *Culicoides* used during the study. PM and KD also wish to acknowledge support funding from EU H20:20 grant PALE-Blu number 727393.

---

**Table S1:**
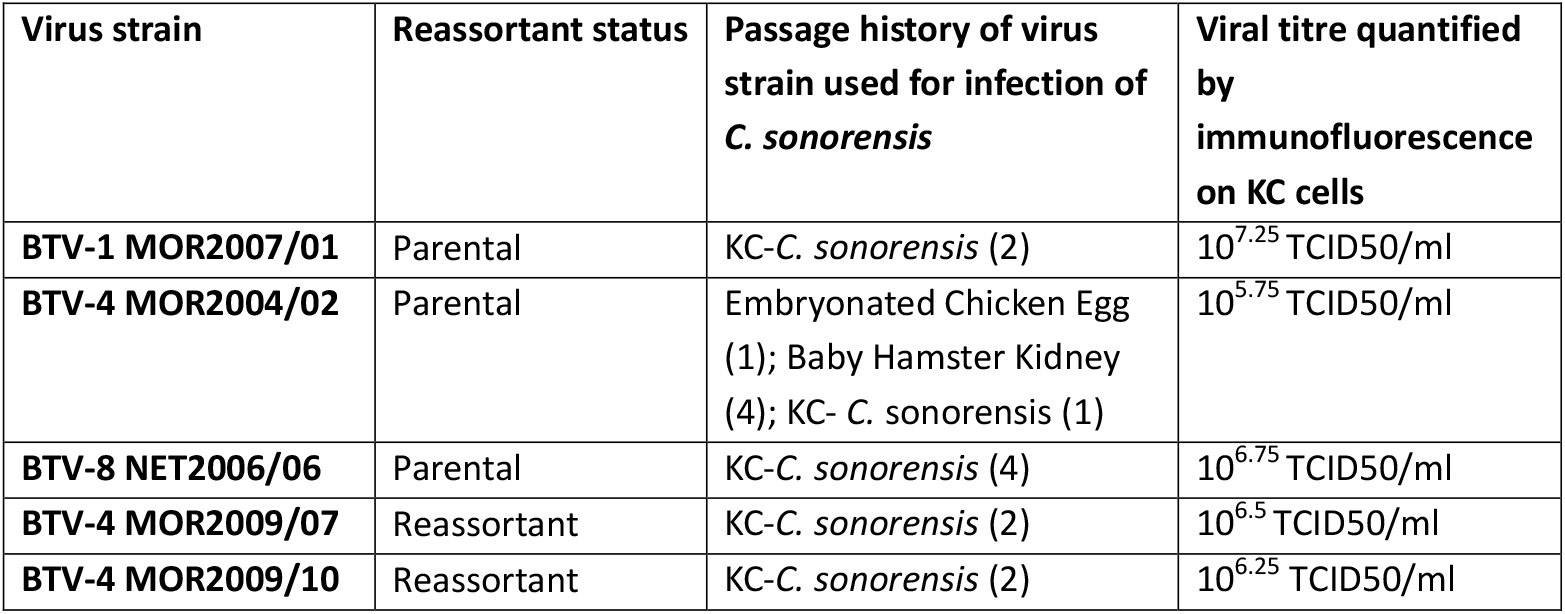
Passage history of virus strains used during studies. All viruses were subjected to a single passage in KC-*C. sonorensis* prior to measurement of titre.

**Table S2.**
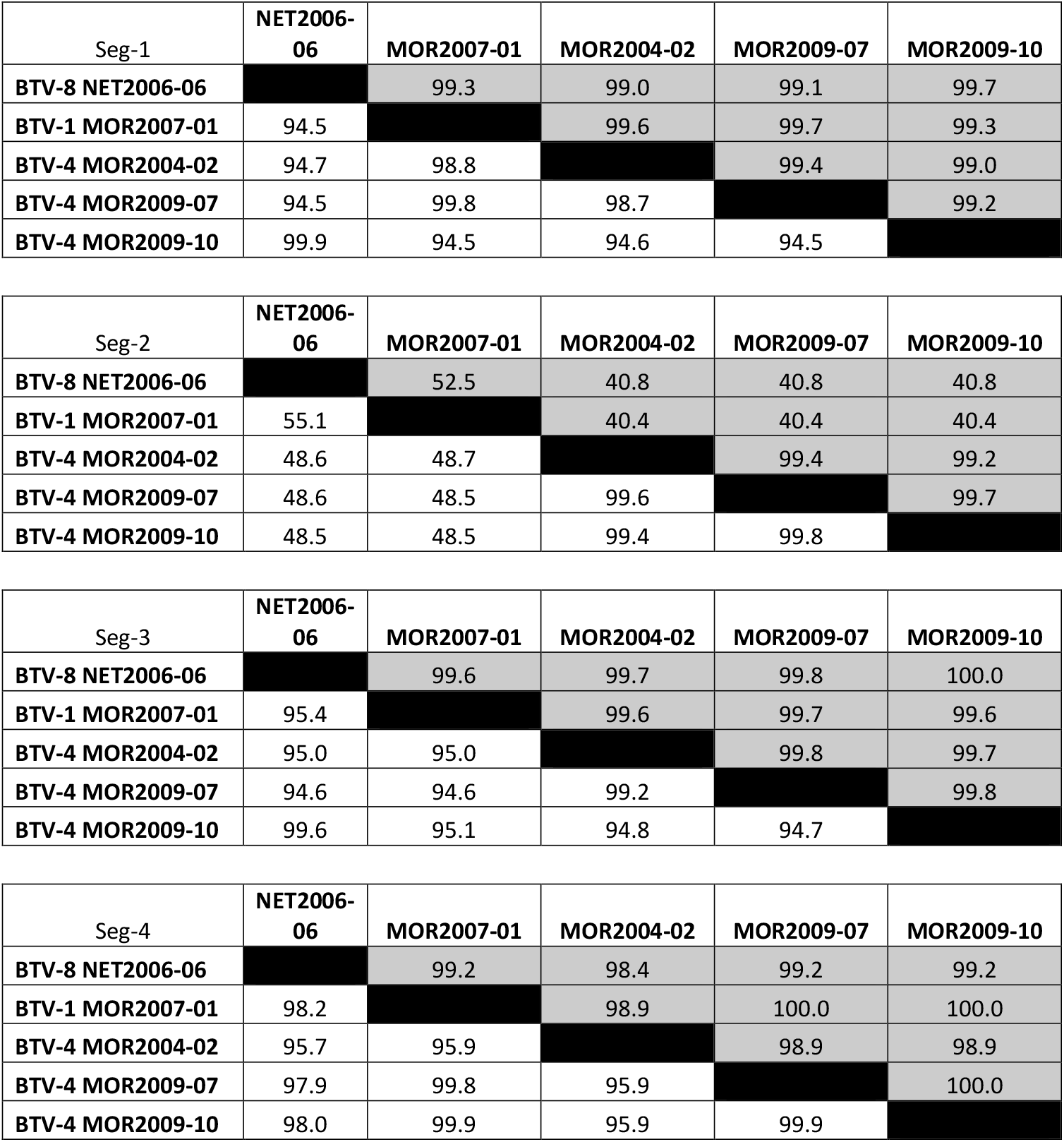
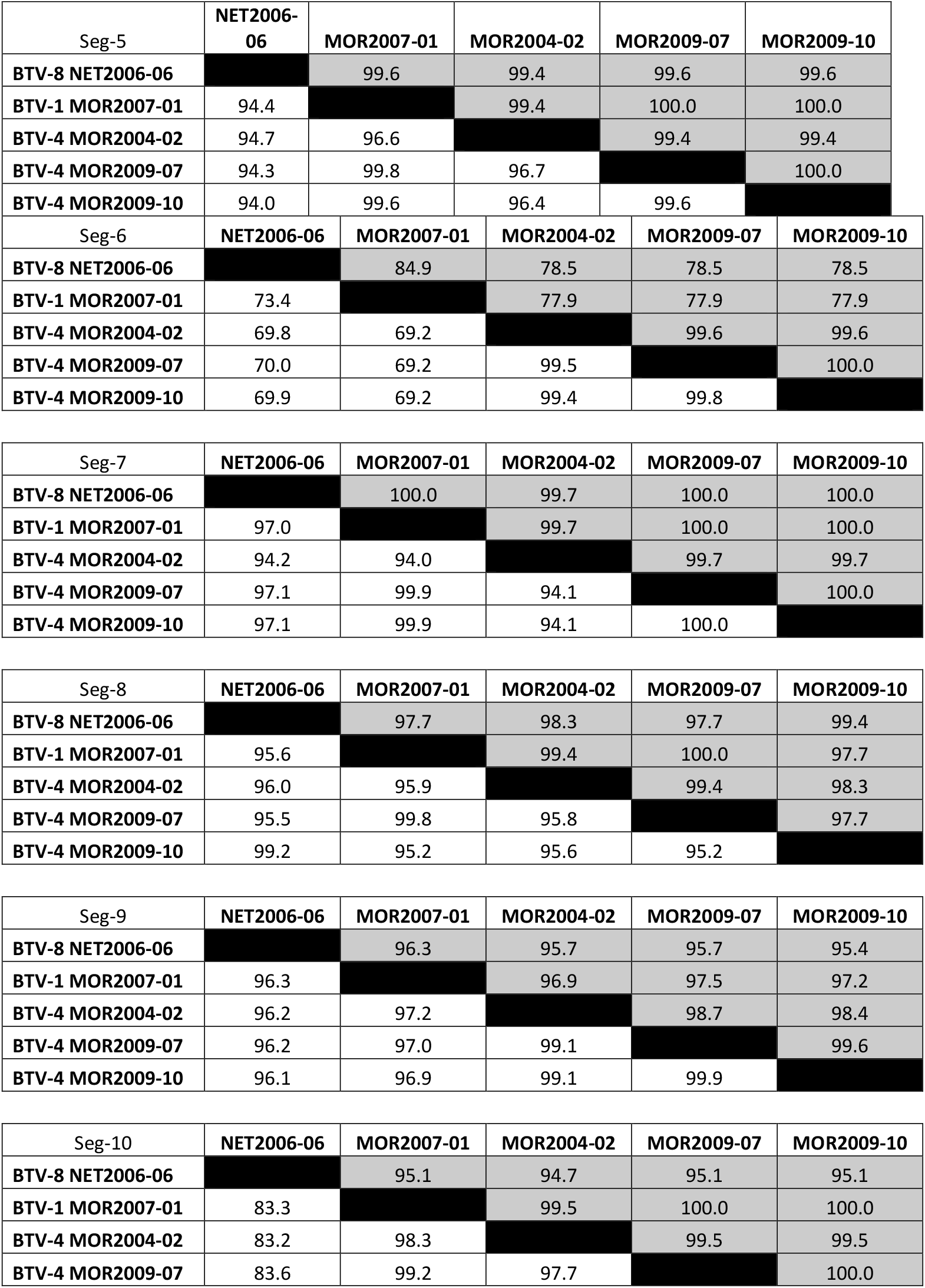
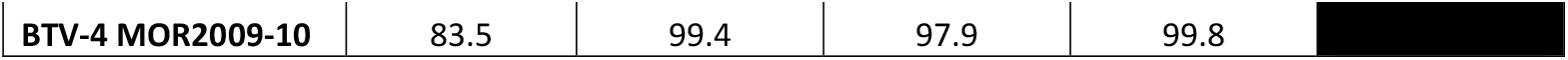
Nucleotide identity (white) and amino acid similarity (grey) expressed as % between bluetongue virus (BTV) strains across all ten genome segments.

**Table S3:**
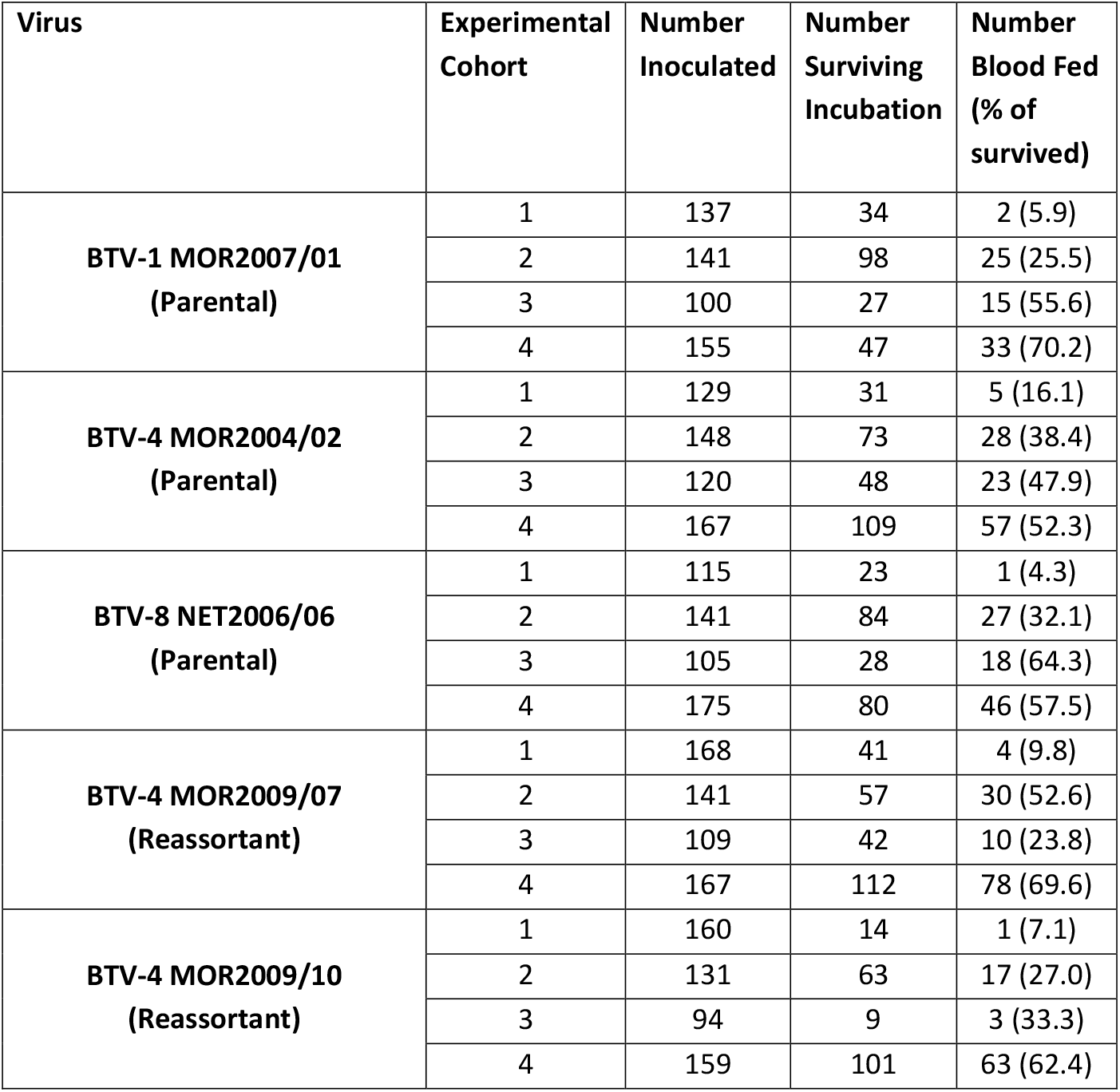
Inoculation and feeding success of *Culicoides sonorensis* infected with five strains of BTV across four cohorts of sheep infection studies.

**Table S4:**
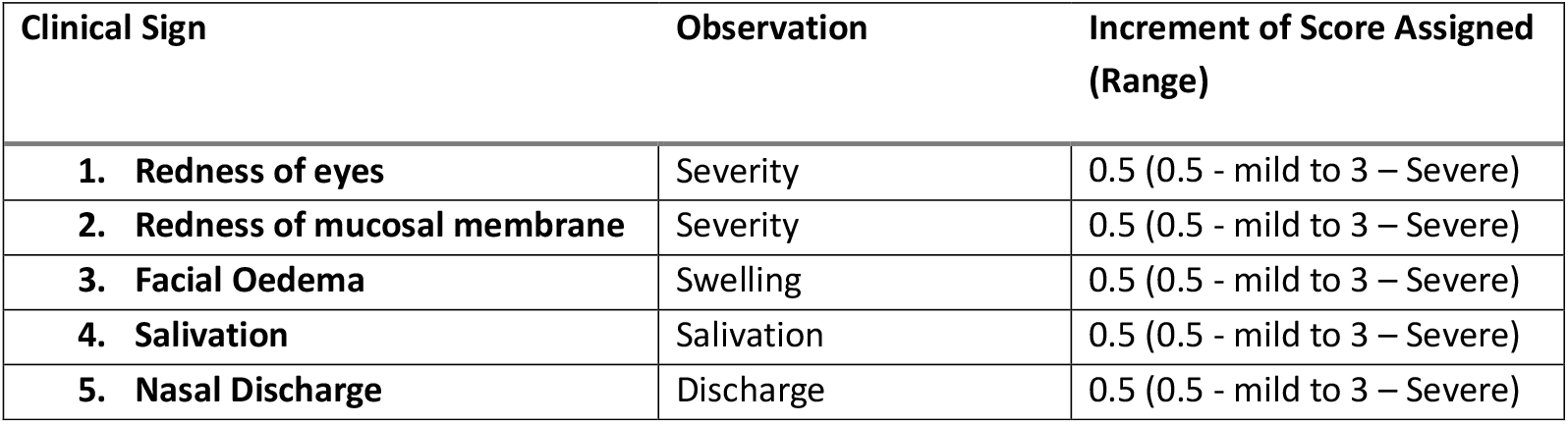
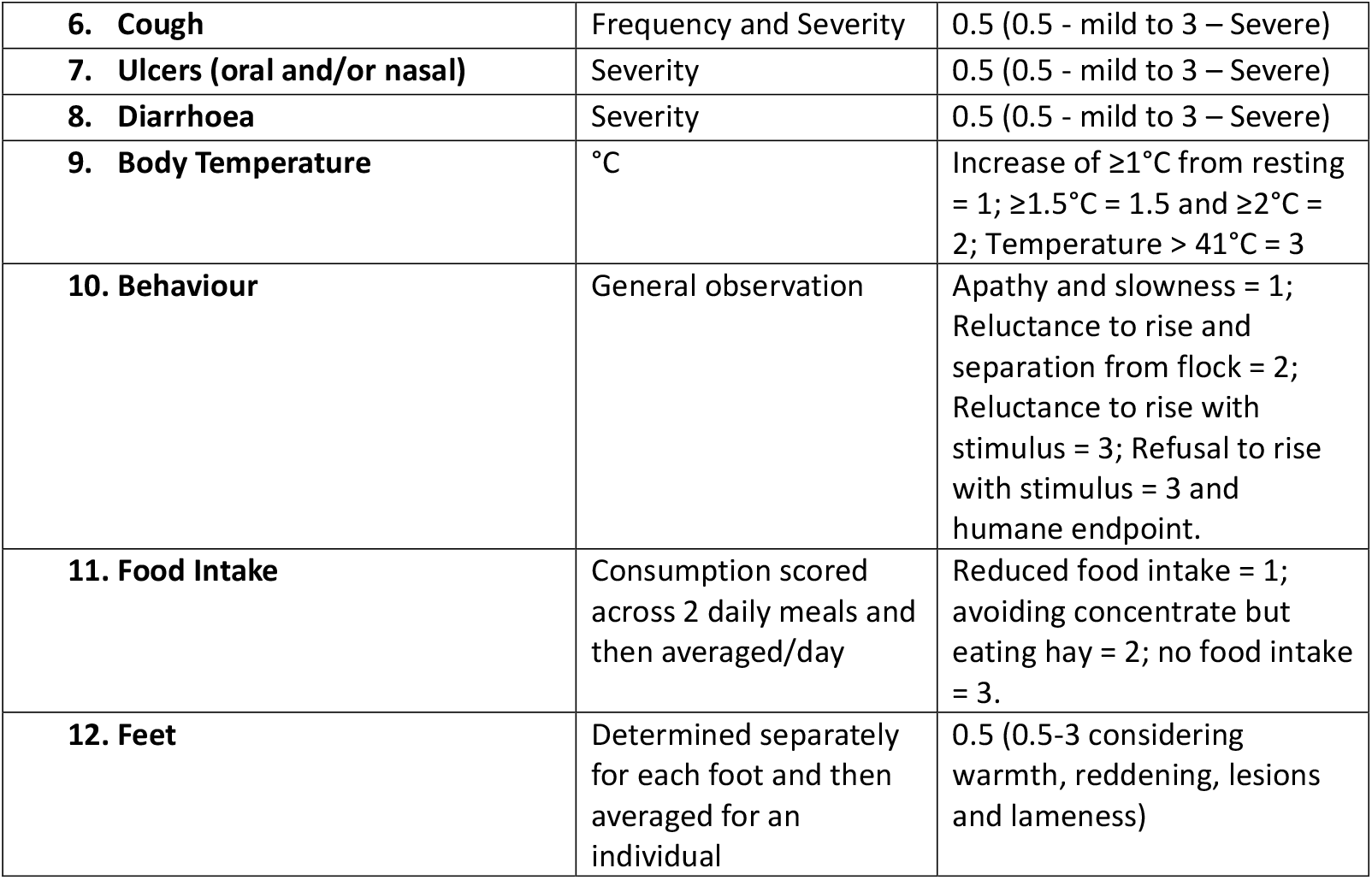
Clinical scoring used to assess severity of disease signs in sheep infected with BTV parental and reassortant strains.

**Table S5:**
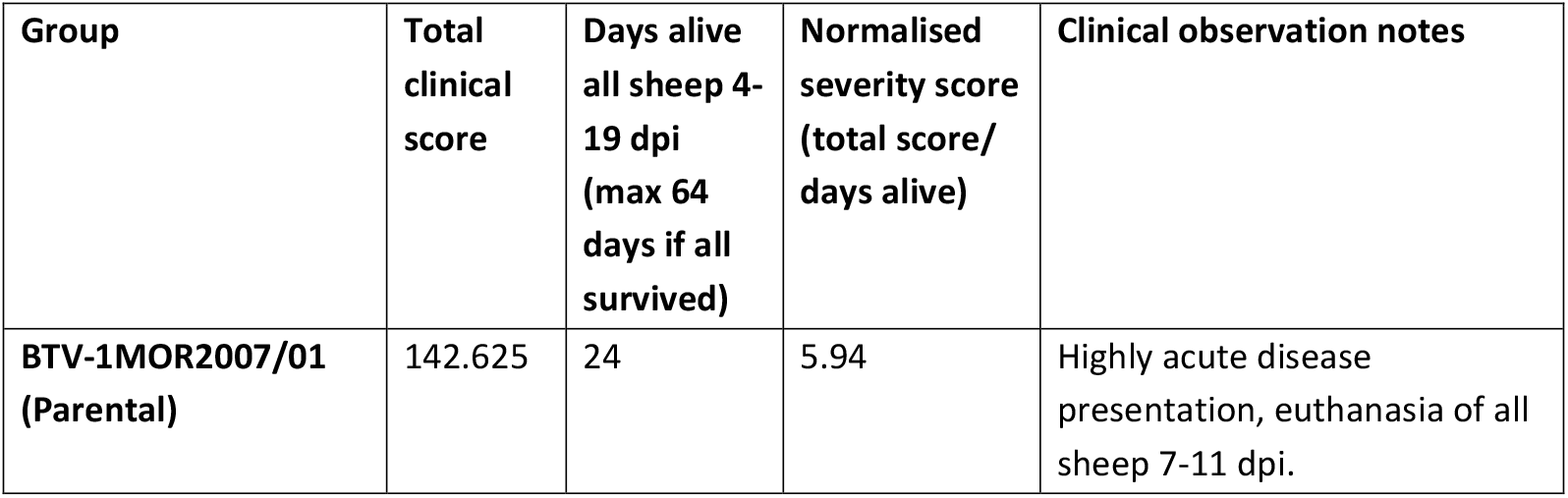
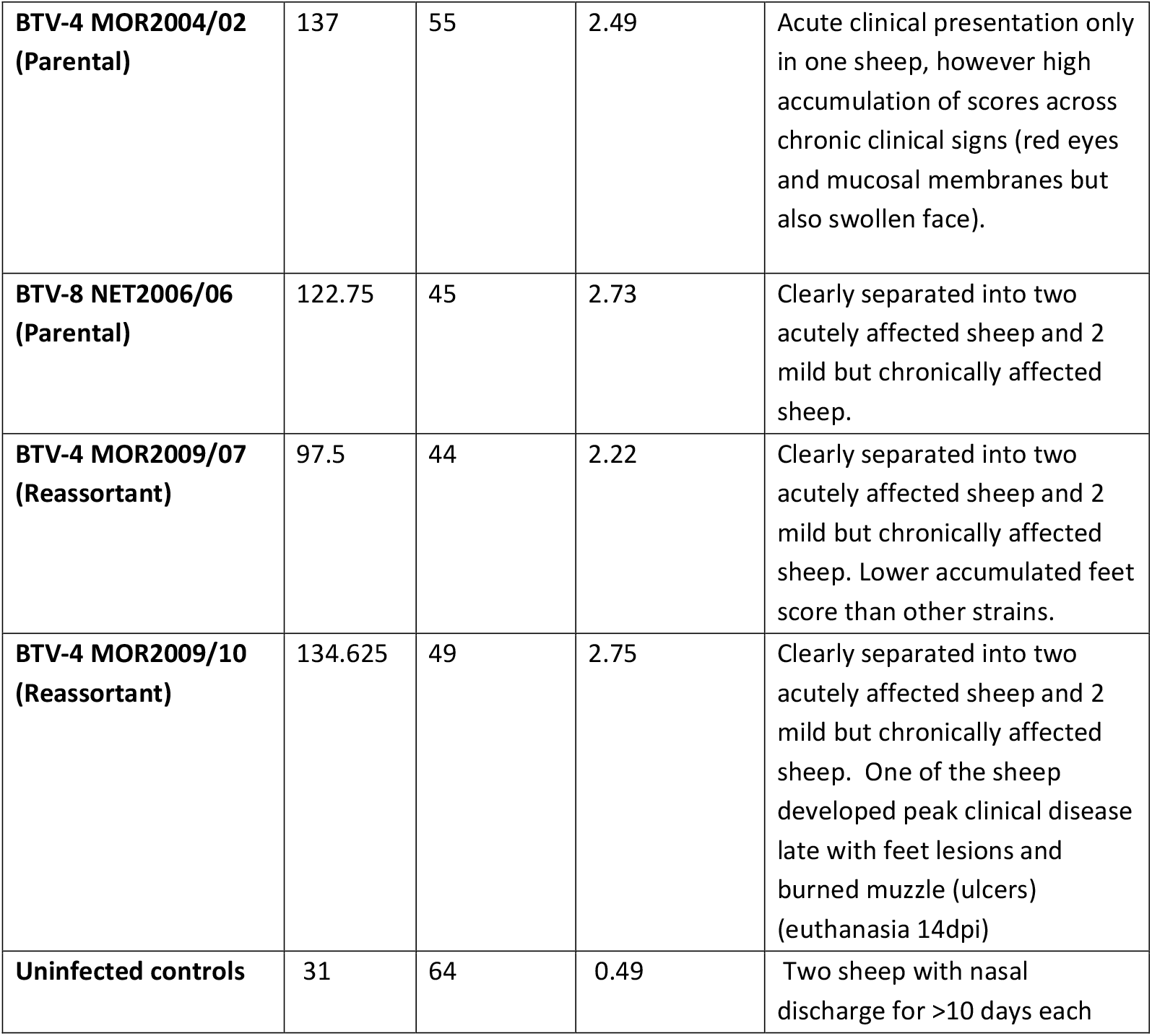
Summary of clinical severity of all five parental and reassortant BTV strains in sheep.

**Table S6.**
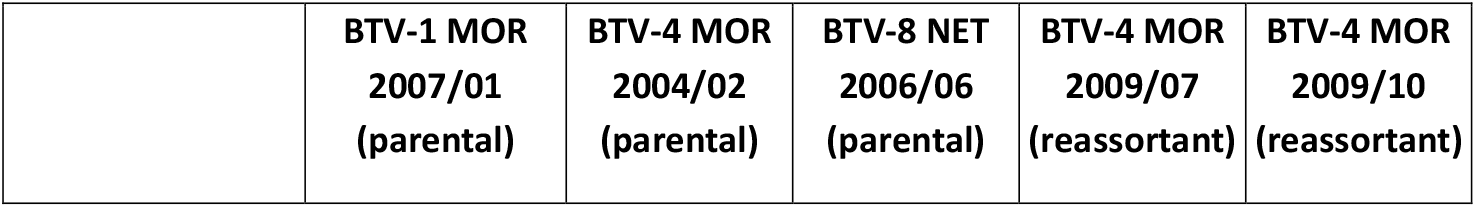
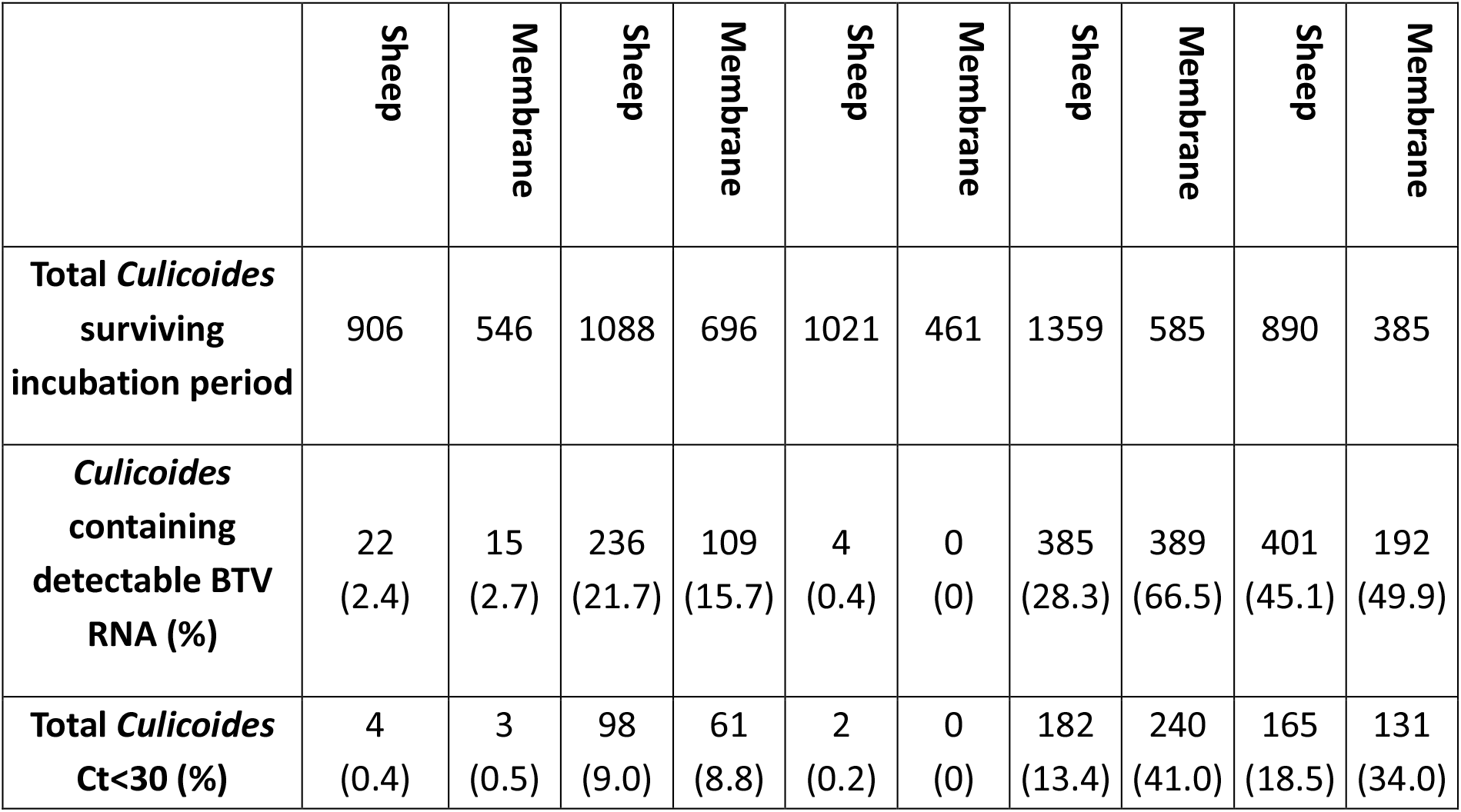
Infection rates of *C. sonorensis* fed either directly on viraemic sheep or indirectly on sheep blood using a membrane-based device.

**Table S7.**
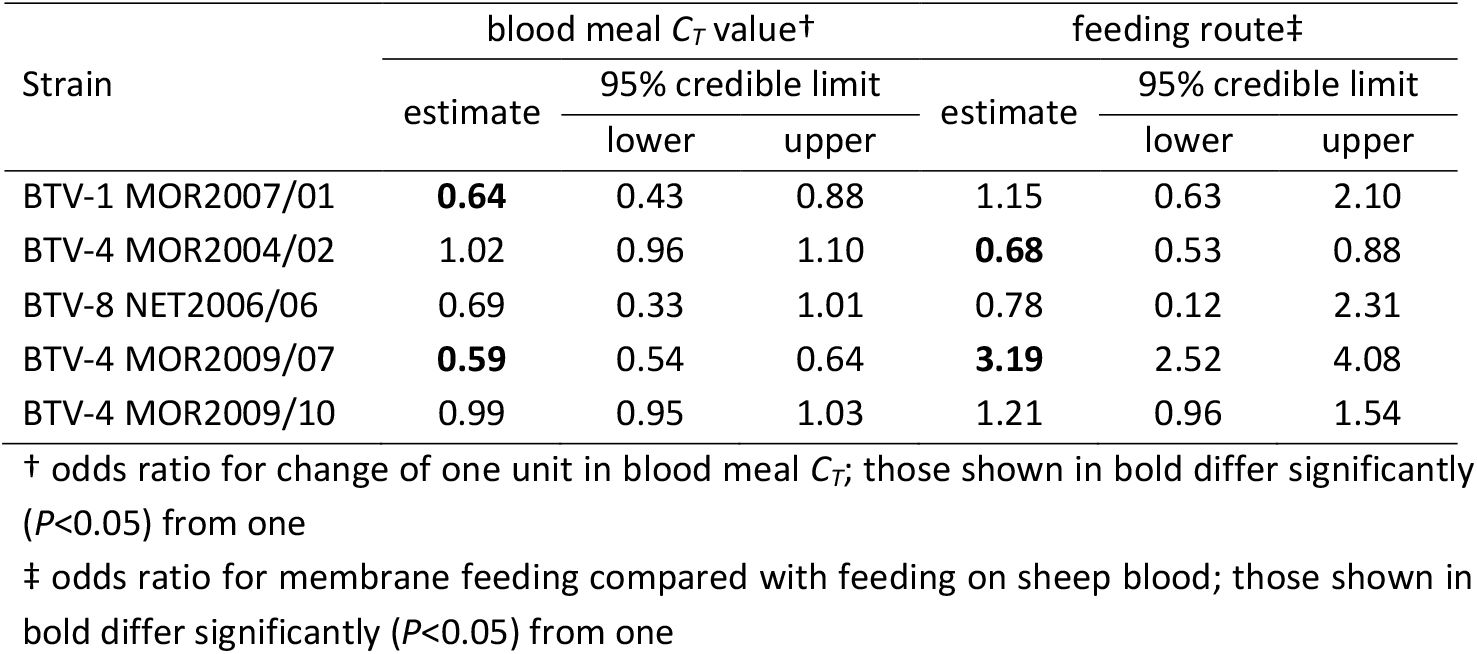
Effect of blood meal *C*_*T*_ value and feeding route on viral infection of *C. sonorensis* for five strains of bluetongue virus.

**Figure S1.**
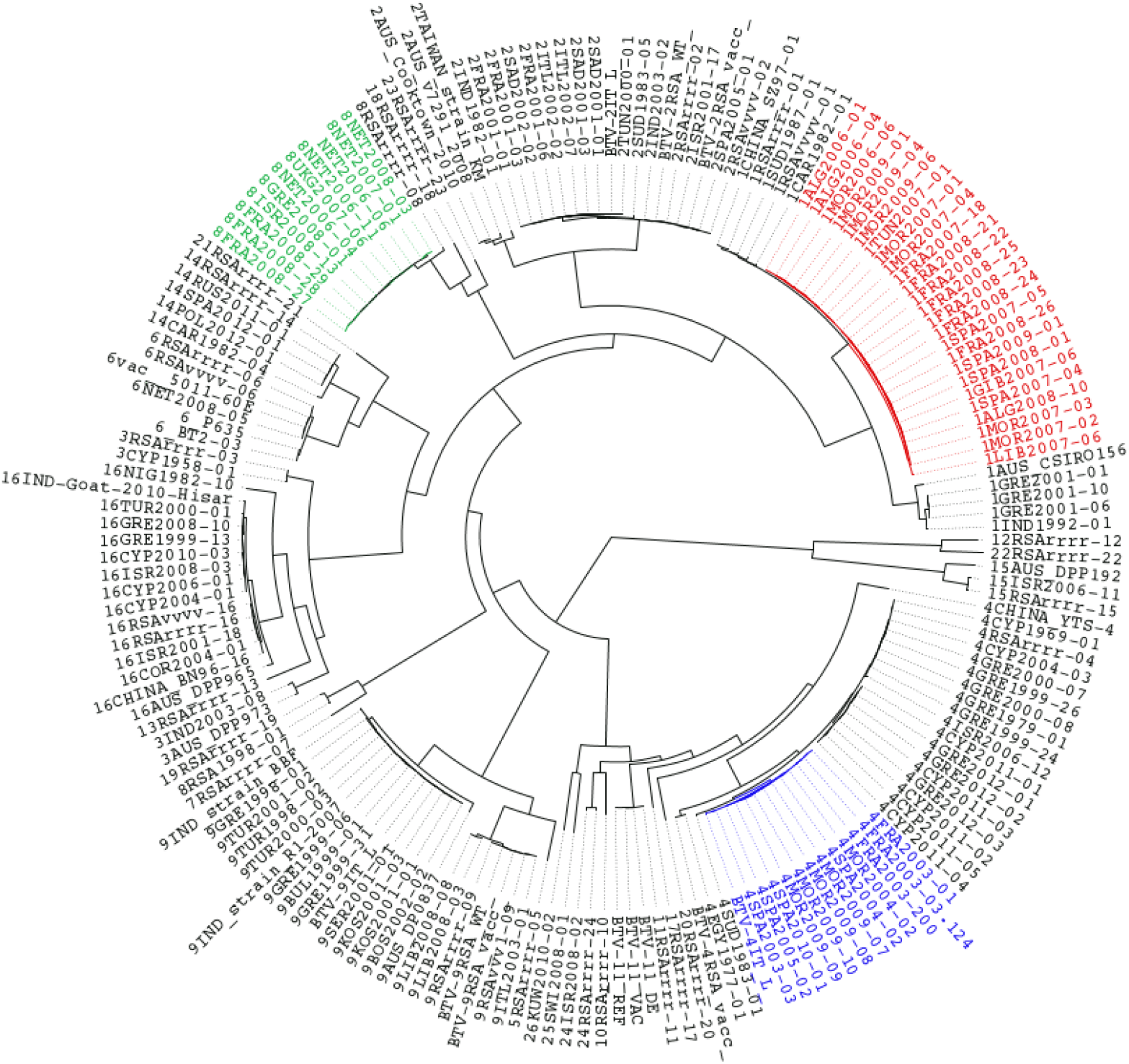
Maximum likelihood tree was constructed for the VP2 coding regions of BTV using IQ-Tree software version 1.3.11.1 (Nguyen, L.T., et al) and the reliability of each tree was estimated by ultrafast bootstrap (Minh, B.Q) analysis of 1000 replicates. The GTR + I + G4 model of evolution was selected according to the Bayesian information criterion score calculated using the IQ-Tree software. Phylogenetic trees were visualised and rooted on the midpoint using the Figtree v1.4.4 software. BTV-1 lineage 2006-2009 is shown in red, BTV-4 lineage 2003-2010 in blue and BTV-8 lineage 2006-2008 in green.

**Figure S2:**
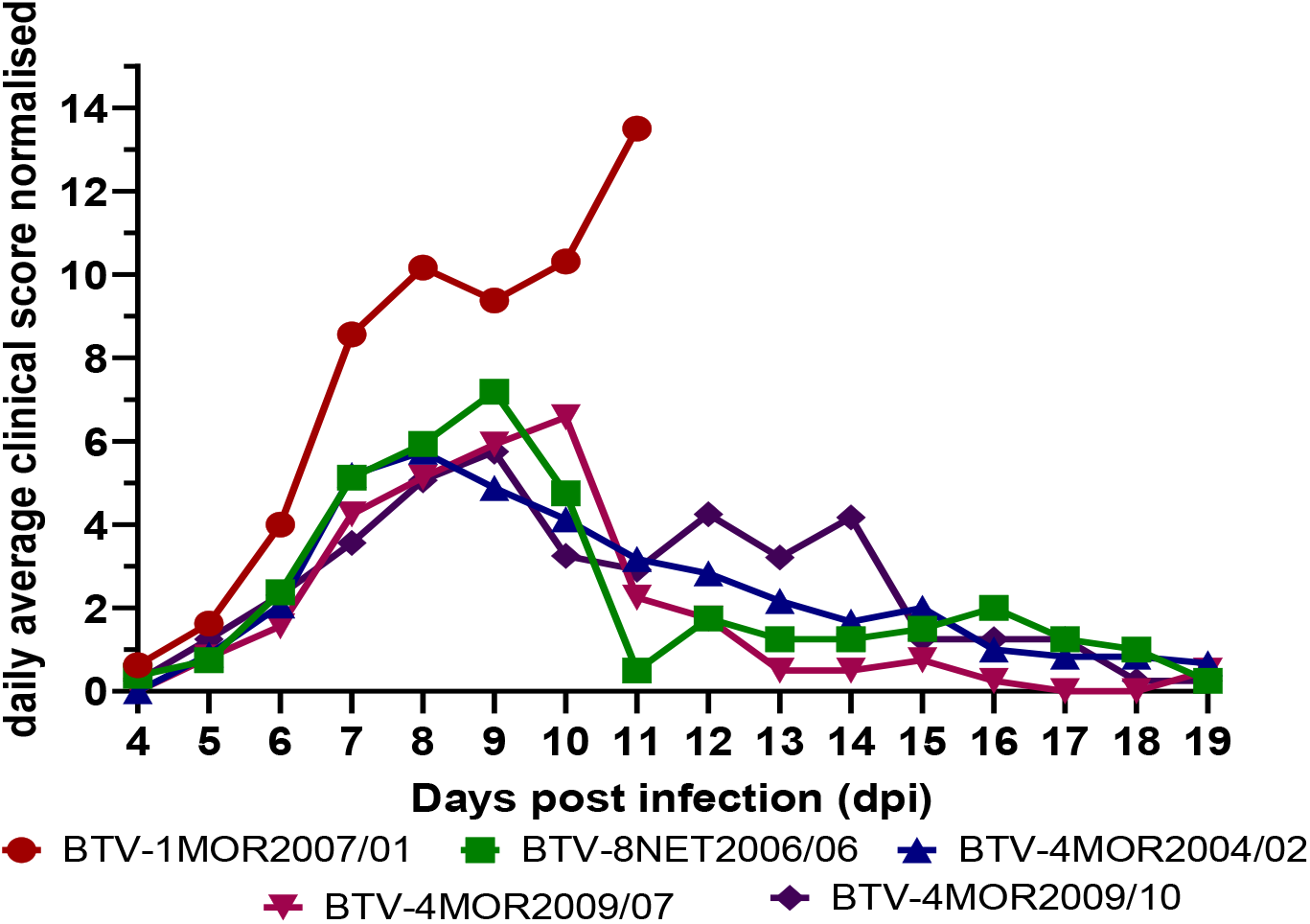
Clinical severity of 5 different BTV strains over time. The daily average normalised clinical score combined for all clinical signs visualised for the clinical period between 4-19 d.p.i. for each of the 5 different BTV strains.

**Figure S3:**
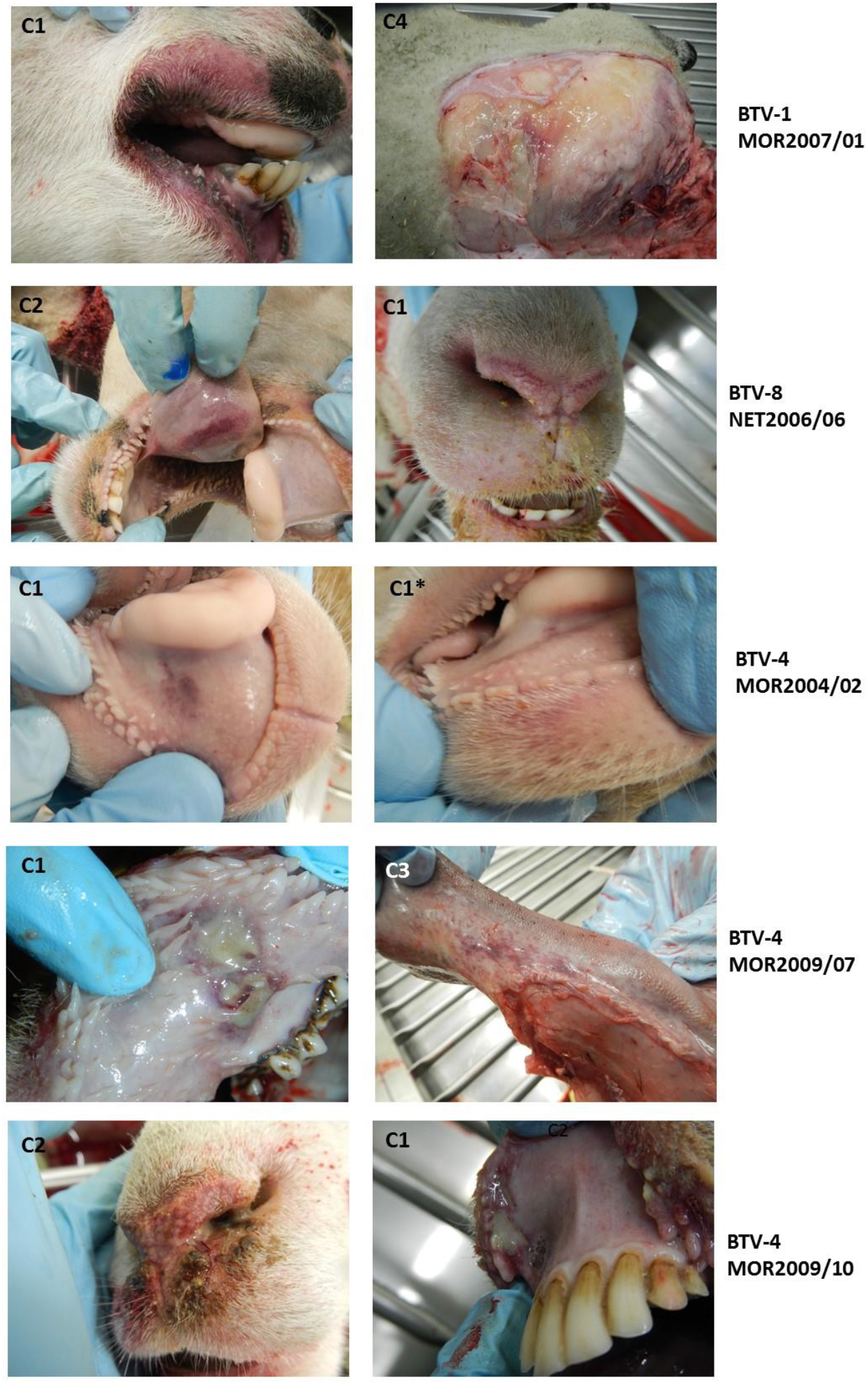
Typical pathological changes in acutely affected sheep infected with 5 different strains of BTV Pathological signs observed in sheep euthanized for reaching humane endpoints between 7-14 days post infection. The cohort of the sheep is given in each panel. BTV-1 MOR2007/01 caused wide ranging haemorrhages in the oral and nasal mucosa, but also skeletal muscles, as well as significant subcutaneous facial and systemic oedema (4/4 sheep). BTV-8 NET2006/06 caused moderate haemorrhagic lesions to the oral and nasal mucosa and tongue (2/4 sheep). BTV4 MOR2004/02 only caused mild haemorrhagic lesions of the oral mucosa (1/4 sheep – both pictures of the same sheep). Both BTV-4 reassortant strains (BTV4 MOR 2009/07 and BTV-4 MOR 2009/10) caused oral and nasal haemorrhagic lesions exceeding those of the BTV-4 and BTV-8 parental strains, but less severe than the BTV-1 parental strain (2/4 sheep for each strain). These two viral strains caused significant oral or nasal ulcerations in the affected sheep that were more severe than those seen for any parental BTV strain.

## REFERENCES

1. McDonald SM, Nelson MI, Turner PE, Patton JT. 2016. Reassortment in segmented RNA viruses: mechanisms and outcomes. Nature Reviews Microbiology 14:448–460.

2. Lakdawala SS, Lamirande EW, Suguitan AL, Wang WJ, Santos CP, Vogel L, Matsuoka Y, Lindsley WG, Jin H, Subbarao K. 2011. Eurasian-Origin Gene Segments Contribute to the Transmissibility, Aerosol Release, and Morphology of the 2009 Pandemic H1N1 Influenza Virus. Plos Pathogens 7.

3. Shelton H, Smith M, Hartgroves L, Stilwell P, Roberts K, Johnson B, Barclay W. 2012. An influenza reassortant with polymerase of pH1N1 and NS gene of H3N2 influenza A virus is attenuated in vivo. Journal of General Virology 93:998–1006.

4. Song MS, Pascua PNQ, Lee JH, Baek YH, Park KJ, Kwon HI, Park SJ, Kim CJ, Kim H, Webby RJ, Webster RG, Choi YK. 2011. Virulence and Genetic Compatibility of Polymerase Reassortant Viruses Derived from the Pandemic (H1N1) 2009 Influenza Virus and Circulating Influenza A Viruses. Journal of Virology 85:6275–6286.

5. Batten CA, Maan S, Shaw AE, Maan NS, Mertens PPC. 2008. A European field strain of bluetongue virus derived from two parental vaccine strains by genome segment reassortment. Virus Research 137:56–63.

6. Nomikou K, Hughes J, Wash R, Kellam P, Breard E, Zientara S, Palmarini M, Biek R, Mertens P. 2015. Widespread Reassortment Shapes the Evolution and Epidemiology of Bluetongue Virus following European Invasion. Plos Pathogens 11.

7. Maclachlan NJ, Drew CP, Darpel KE, Worwa G. 2009. The Pathology and Pathogenesis of Bluetongue. Journal of Comparative Pathology 141:1–16.

8. Purse BV, Carpenter S, Venter GJ, Bellis G, Mullens BA. 2015. Bionomics of Temperate and Tropical Culicoides Midges: Knowledge Gaps and Consequences for Transmission of Culicoides-Borne Viruses, p 373-+. In Berenbaum MR (ed), Annual Review of Entomology, Vol 60, vol 60.

9. Carpenter S, Veronesi E, Mullens B, Venter G. 2015. Vector competence of Culicoides for arboviruses: three major periods of research, their influence on current studies and future directions. Revue Scientifique Et Technique-Office International Des Epizooties 34:97–112.

10. Darpel KE, Batten C, Veronesi E, Shaw A, Anthony S, Bachanek-Bankowska K, Kgosana L, Bin-Tarif A, Carpenter S, Mueller-Doblies U, Takamatsu H-H, Mellor PS, Mertens PP, Oura CAL. 2012. Clinical signs and pathology shown by British sheep and cattle infected with bluetongue virus serotype 8 derived from the 2006 outbreak in northern Europe Veterinary Record 161:253–261.

11. Melzi E, Caporale M, Rocchi M, Martin V, Gamino V, di Provvido A, Marruchella G, Entrican G, Sevilla N, Palmarini M. 2016. Follicular dendritic cell disruption as a novel mechanism of virus-induced immunosuppression. Proceedings of the National Academy of Sciences of the United States of America 113:E6238–E6247.

12. Elbers ARW, Backx A, Meroc E, Gerbier G, Staubach C, Hendrickx G, van der Spek A, Mintiens K. 2008. Field observations during the bluetongue serotype 8 epidemic in 2006 - I. Detection of first outbreaks and clinical signs in sheep and cattle in Belgium, France and the Netherlands. Preventive Veterinary Medicine 87:21–30.

13. Elbers ARW, Backx A, Mintiens K, Gerbier G, Staubach C, Hendrickx G, van der Spek A. 2008. Field observations during the Bluetongue serotype 8 epidemic in 2006 - II. Morbidity and mortality rate, case fatality and clinical recovery in sheep and cattle in the Netherlands. Preventive Veterinary Medicine 87:31–40.

14. Purse BV, Mellor PS, Rogers DJ, Samuel AR, Mertens PPC, Baylis M. 2005. Climate change and the recent emergence of bluetongue in Europe. Nature Reviews Microbiology 3:171–181.

15. Carpenter S, Wilson A, Mellor PS. 2009. Culicoides and the emergence of bluetongue virus in northern Europe. Trends in Microbiology 17:172–178.

16. Flannery J, Sanz-Bernardo B, Ashby M, Brown H, Carpenter S, Cooke L, Corla A, Frost L, Gubbins S, Hicks H, Qureshi M, Rajko-Nenow P, Sanders C, Tully M, Breard E, Sailleau C, Zientara S, Darpel K, Batten C. 2019. Evidence of reduced viremia, pathogenicity and vector competence in a re-emerging European strain of bluetongue virus serotype 8 in sheep. Transboundary and Emerging Diseases 66:1177–1185.

17. Lorusso A, Sghaier S, Carvelli A, Di Gennaro A, Leone A, Marini V, Pelini S, Marcacci M, Rocchigiani AM, Puggioni G, Savini G. 2013. Bluetongue virus serotypes 1 and 4 in Sardinia during autumn 2012: New incursions or re-infection with old strainsã Infection Genetics and Evolution 19:81–87.

18. Shaw AE, Ratinier M, Nunes SF, Nomikou K, Caporale M, Golder M, Allan K, Hamers C, Hudelet P, Zientara S, Breard E, Mertens P, Palmarini M. 2013. Reassortment between Two Serologically Unrelated Bluetongue Virus Strains Is Flexible and Can Involve any Genome Segment. Journal of Virology 87:543–557.

19. Ratinier M, Caporale M, Golder M, Franzoni G, Allan K, Nunes SF, Armezzani A, Bayoumy A, Rixon F, Shaw A, Palmarini M. 2011. Identification and Characterization of a Novel Non-Structural Protein of Bluetongue Virus. Plos Pathogens 7.

20. Roy P, Noad R. 2006. Bluetongue virus assembly and morphogenesis. Reoviruses: Entry, Assembly and Morphogenesis 309:87–116.

21. Oberst RD, Stott JL, Blanchardchannell M, Osburn BI. 1987. Genetic reassortment of bluetongue virus serotype-11 strains in the bovine. Veterinary Microbiology 15:11–18.

22. Samal SK, Elhussein A, Holbrook FR, Beaty BJ, Ramig RF. 1987. Mixed infection off Culicoides variipennis with bluetongue virus serotype 10 and serotype 17 - evidence for high-frequency reassortment in the vector. Journal of General Virology 68:2319–2329.

23. Samal SK, Livingston CW, McConnell S, Ramig RF. 1987. Analysis of mixed infection of sheep with bluetongue virus serotype 10 and serotype 17 - evidence for genetic reassortment in the vertebrate host. Journal of Virology 61:1086–1091.

24. Stott JL, Oberst RD, Channell MB, Osburn BI. 1987. Genome segement reassortment between 2 serotypes of bluetongue virus in a natural host. Journal of Virology 61:2670–2674.

25. Wittmann EJ, Mellor PS, Baylis M. 2002. Effect of temperature on the transmission of orbiviruses by the biting midge, Culicoides sonorensis. Medical and Veterinary Entomology 16:147–156.

26. Riegler. 2002. Variation in African horse sickness virus and its effect on the vector competence of Culicoides biting midges. PhD ThesisUniversity of Surrey.

27. Carpenter S, Lunt HL, Arav D, Venter GJ, Mellor PS. 2006. Oral susceptibility to bluetongue virus of Culicoides (Diptera : Ceratopogonidae) from the United Kingdom. Journal of Medical Entomology 43:73–78.

28. Tabachnick WJ. 1991. Genetic control of oral-susceptibility to infection of Culicoides variipennnis with bluetongue virus. American Journal of Tropical Medicine and Hygiene 45:666–671.

29. Brenner J, Batten C, Yadin H, Bumbarov V, Friedgut O, Rotenberg D, Golender N, Oura CAL. 2011. Clinical syndromes associated with the circulation of multiple serotypes of bluetongue virus in dairy cattle in Israel. Veterinary Record 169:389–U41.

30. Dal Pozzo F, Martinelle L, Thys C, Sarradin P, De Leeuw I, Van Campe W, De Clercq K, Thiry E, Saegerman C. 2013. Experimental co-infections of calves with bluetongue virus serotypes 1 and 8. Veterinary Microbiology 165:167–172.

31. Sanchez-Cordon PJ, Pleguezuelos FJ, de Diego ACP, Gomez-Villamandos JC, Sanchez-Vizcaino JM, Ceron JJ, Tecles F, Garfia B, Pedrera M. 2013. Comparative study of clinical courses, gross lesions, acute phase response and coagulation disorders in sheep inoculated with bluetongue virus serotype 1 and 8. Veterinary Microbiology 166:184–194.

32. Caporale M, Wash R, Pini A, Savini G, Franchi P, Golder M, Patterson-Kane J, Mertens P, Di Gialleonardo L, Armillotta G, Lelli R, Kellam P, Palmarini M. 2011. Determinants of Bluetongue Virus Virulence in Murine Models of Disease. Journal of Virology 85:11479–11489.

33. Celma CC, Bhattacharya B, Eschbaumer M, Wernike K, Beer M, Roy P. 2014. Pathogenicity study in sheep using reverse-genetics-based reassortant bluetongue viruses. Veterinary Microbiology 174:139–147.

34. Janowicz A, Caporale M, Shaw A, Gulletta S, Di Gialleonardo L, Ratinier M, Palmarini M. 2015. Multiple Genome Segments Determine Virulence of Bluetongue Virus Serotype 8. Journal of Virology 89:5238–5249.

35. Coetzee P, Van Vuuren M, Stokstad M, Myrmel M, Venter EH. 2012. Bluetongue virus genetic and phenotypic diversity: Towards identifying the molecular determinants that influence virulence and transmission potential. Veterinary Microbiology 161:1–12.

36. Coetzee P, van Vuuren M, Venter EH, Stokstad M. 2014. A review of experimental infections with bluetongue virus in the mammalian host. Virus Research 182:21–34.

37. van Gennip RGP, Drolet BS, Lopez PR, Roost AJC, Boonstra J, van Rijn PA. 2019. Vector competence is strongly affected by a small deletion or point mutations in bluetongue virus. Parasites & Vectors 12:16.

38. Jeggo MH, Gumm ID, Taylor WP. 1983. Clinical and serological response of sheep to serial challenge with different bluetongue virus types. Research in Veterinary Science 34:205–211.

39. Jones RH, Foster NM. 1978. Heterogeneity of Culicoides variipennis field populations to oral infection with bluetongue virus. American Journal of Tropical Medicine and Hygiene 27:178–183.

40. Veronesi E, Darpel K, Gubbins S, Batten C, Nomikou K, Mertens P, Carpenter S. 2020. Diversity of Transmission Outcomes Following Co-Infection of Sheep with Strains of Bluetongue Virus Serotype 1 and 8. Microorganisms 8:16.

41. Baylis M, O’Connell L, Mellor PS. 2008. Rates of bluetongue virus transmission between Culicoides sonorensis and sheep. Medical and Veterinary Entomology 22:228–237.

42. van Wuijckhuise L, Dercksen D, Muskens J, de Bruyn J, Scheepers M, Vrouenraets R. 2006. Bluetongue in the Netherlands; description of the first clinical cases and differential diagnosis; Common symptoms just a little different and in too many herds. Tijdschrift Voor Diergeneeskunde 131:649–654.

43. Sanders CJ, Shortall CR, Gubbins S, Burgin L, Gloster J, Harrington R, Reynolds DR, Mellor PS, Carpenter S. 2011. Influence of season and meteorological parameters on flight activity of Culicoides biting midges. Journal of Applied Ecology 48:1355–1364.

44. Searle KR, Barber J, Stubbins F, Labuschagne K, Carpenter S, Butler A, Denison E, Sanders C, Mellor PS, Wilson A, Nelson N, Gubbins S, Purse BV. 2014. Environmental Drivers of Culicoides Phenology: How Important Is Species-Specific Variation When Determining Disease Policyã Plos One 9:13.

45. Darpel KE, Monaghan P, Simpson J, Anthony SJ, Veronesi E, Brooks HW, Elliott H, Brownlie J, Takamatsu H-H, Mellor PS, Mertens PPC. 2012. Involvement of the skin during bluetongue virus infection and replication in the ruminant host. Veterinary Research 43.

46. Busquets MG, Pullinger GD, Darpel KE, Cooke L, Armstrong S, Simpson J, Palmarini M, Fragkoudis R, Mertens PPC. 2021. An Early Block in the Replication of the Atypical Bluetongue Virus Serotype 26 in Culicoides Cells Is Determined by Its Capsid Proteins. Viruses-Basel 13.

47. Pullinger GD, Busquets MG, Nomikou K, Boyce M, Attoui H, Mertens PP. 2016. Identification of the Genome Segments of Bluetongue Virus Serotype 26 (Isolate KUW2010/02) that Restrict Replication in a Culicoides sonorensis Cell Line (KC Cells). Plos One 11.

48. Hassan SS, Roy P. 1999. Expression and functional characterization of bluetongue virus VP2 protein: Role in cell entry. Journal of Virology 73:9832–9842.

49. Hassan SH, Wirblich C, Forzan M, Roy P. 2001. Expression and functional characterization of bluetongue virus VP5 protein: Role in cellular permeabilization. Journal of Virology 75:8356–8367.

50. Darpel KE, Langner KFA, Nimtz M, Anthony SJ, Brownlie J, Takamatsu H-H, Mellor PS, Mertens PPC. 2011. Saliva Proteins of Vector Culicoides Modify Structure and Infectivity of Bluetongue Virus Particles. Plos One 6.

51. Fu H, Leake CJ, Mertens PPC, Mellor PS. 1999. The barriers to bluetongue virus infection, dissemination and transmission in the vector, Culicoides variipennis (Diptera : Ceratopogonidae). Archives of Virology 144:747–761.

52. Kreuger F. Accessed on 10 December 2019 2009. Trim Galore.https://githubcom/FelixKrueger/TrimGalore2019.

53. Li H, Durbin R. 2009. Fast and accurate short read alignment with Burrows-Wheeler transform. Bioinformatics 25:1754–1760.

54. Martin DP, Murrell B, Golden M, Khoosal A, Muhire B. 2015. RDP4: Detection and analysis of recombination patterns in virus genomes. Virus Evolution 1:5.

55. Tamura K, Stecher G, Peterson D, Filipski A, Kumar S. 2013. MEGA6: Molecular Evolutionary Genetics Analysis Version 6.0. Molecular Biology and Evolution 30:2725–2729.

56. Vaidya G, Lohman DJ, Meier R. 2011. SequenceMatrix: concatenation software for the fast assembly of multi-gene datasets with character set and codon information. Cladistics 27:171–180.

57. Maan S, Maan NS, Belaganahalli MN, Potgieter AC, Kumar V, Batra K, Wright IM, Kirkland PD, Mertens PPC. 2016. Development and Evaluation of Real Time RT-PCR Assays for Detection and Typing of Bluetongue Virus. Plos One 11:19.

58. Nayduch D, Cohnstaedt LW, Saski C, Lawson D, Kersey P, Fife M, Carpenter S. 2014. Studying Culicoides vectors of BTV in the post-genomic era: Resources, bottlenecks to progress and future directions. Virus Research 182:43–49.

59. Boorman J. 1974. The maintenance of laboratory colonies of Culicoides variipennis (Coq.), C. nubeculosus (Mg.) and C. riethi Kieff. (Diptera, Ceratopogonidae). Bulletin of Entomological Research 64:371.

60. Veronesi E, Antony F, Gubbins S, Golding N, Blackwell A, Mertens PPC, Brownlie J, Darpel KE, Mellor PS, Carpenter S. 2013. Measurement of the Infection and Dissemination of Bluetongue Virus in Culicoides Biting Midges Using a Semi-Quantitative RT-PCR Assay and Isolation of Infectious Virus. Plos One 8.

61. Shaw AE, Monaghan P, Alpar HO, Anthony S, Darpel KE, Batten CA, Guercio A, Alimena G, Vitale M, Bankowska K, Carpenter S, Jones H, Oura CAL, King DP, Elliott H, Mellor PS, Mertens PPC. 2007. Development and initial evaluation of a real-time RT-PCR assay to detect bluetongue virus genome segment 1. Journal of Virological Methods 145:115–126.

